# Strigolactones optimise plant water usage by modulating vessel formation

**DOI:** 10.1101/2023.04.05.535530

**Authors:** Jiao Zhao, Dongbo Shi, Kiara Kaeufer, Changzheng Song, Xiaocai Xu, Laura Luzzietti, Tom Bennett, Kerstin Kaufmann, Thomas Greb

## Abstract

Wood formation is fundamental for the remarkable growth of plant bodies by continuously providing cells for long-distance transport of water and nutrients^1–3^. Vessel elements, the water transporting units within woody tissues, are generated from a cylindrical domain of cambium stem cells (CSCs) producing different vascular cell types in a bidirectional manner^4–6^. However, knowledge on the regulation of CSC activity is limited with unclear cell fate trajectories as the most obscure aspect in this context. Here, via revealing transcriptome signatures of CSCs and their derivatives with single cell resolution in *Arabidopsis thaliana*, we discover that the strigolactone (SL) signalling pathway modulates cell type composition in vascular tissues and thereby increases drought resistance. In particular, we find that SL signalling negatively regulates vessel element formation and thereby plant water usage. SL signalling is generally associated with differentiating vascular tissues but low in developing vessels and in CSCs implying a local role during fate decisions in CSC-derived cells. Highlighting the importance of vascular tissue composition for the overall plant water balance, alteration of vessel element formation has a direct impact on transpiration rates through leaf stomata. Our results demonstrate the importance of structural alignment of water transporting tissues to unstable water regimes and provide perspectives for a long-term modulation of drought resistance in plants.

## Introduction

Continuous growth and tissue formation is one characteristic of plant development and important for aligning distinct body structures with changing environmental conditions. Cambium-driven radial growth of shoots and roots of dicotyledonous species is an important feature of this growth mode, and key for biomass production and for the long-term sequestration of CO_2_^7^. CSCs proliferate usually providing wood (i.e. xylem) cells inwards and bast (i.e. phloem) cells outwards. Within xylem and phloem tissues, various cell types fulfil highly specialised functions like water transport by xylem vessel elements or sugar transport by phloem sieve elements^8^. However, mechanisms regulating CSC-associated cell fate decisions to cope with stressful environmental conditions are largely unknown.

Here, we provide cell-resolved transcriptomes of CSCs and all CSC-derived cell types generated by single nucleus RNA-sequencing (snRNA-seq). Based on our droplet-and plate-based high-resolution transcriptome analysis, we identify SL signalling as a modulator of vessel formation and, thereby, as a module putatively mediating structural adaptations to fluctuating water availability and drought stress.

## Results and Discussion

### snRNA-seq analysis of radial plant growth

To comprehensively reveal cell states during radial plant growth, we established an snRNA-seq-based atlas of the Arabidopsis hypocotyl, the organ connecting shoot and root systems and being a hotspot for cambium-based organ growth^9^. To this end, nuclei were extracted, filtered, and purified via a cell sorter and processed by droplet-based single nucleus transcriptome analysis^10^ (Fig. 1a, Supplementary Fig. S1a, b). Demonstrating the success of our approach, at least 400 transcribed genes were detected in 4,722 out of 6,780 processed nuclei. From those, we kept 2,061 high quality nuclei in which at least 1,200 unique molecular identifier (UMI) counts were identified for further analysis. Within these high quality nuclei, we detected 26,613 genes as being transcribed, covering nearly 70 % of the annotated genes for Arabidopsis^11^. On the median level, our analysis detected 1,224 transcribed genes and 1,854 UMIs for each nucleus (Supplementary Data 1).

**Fig. 1:**
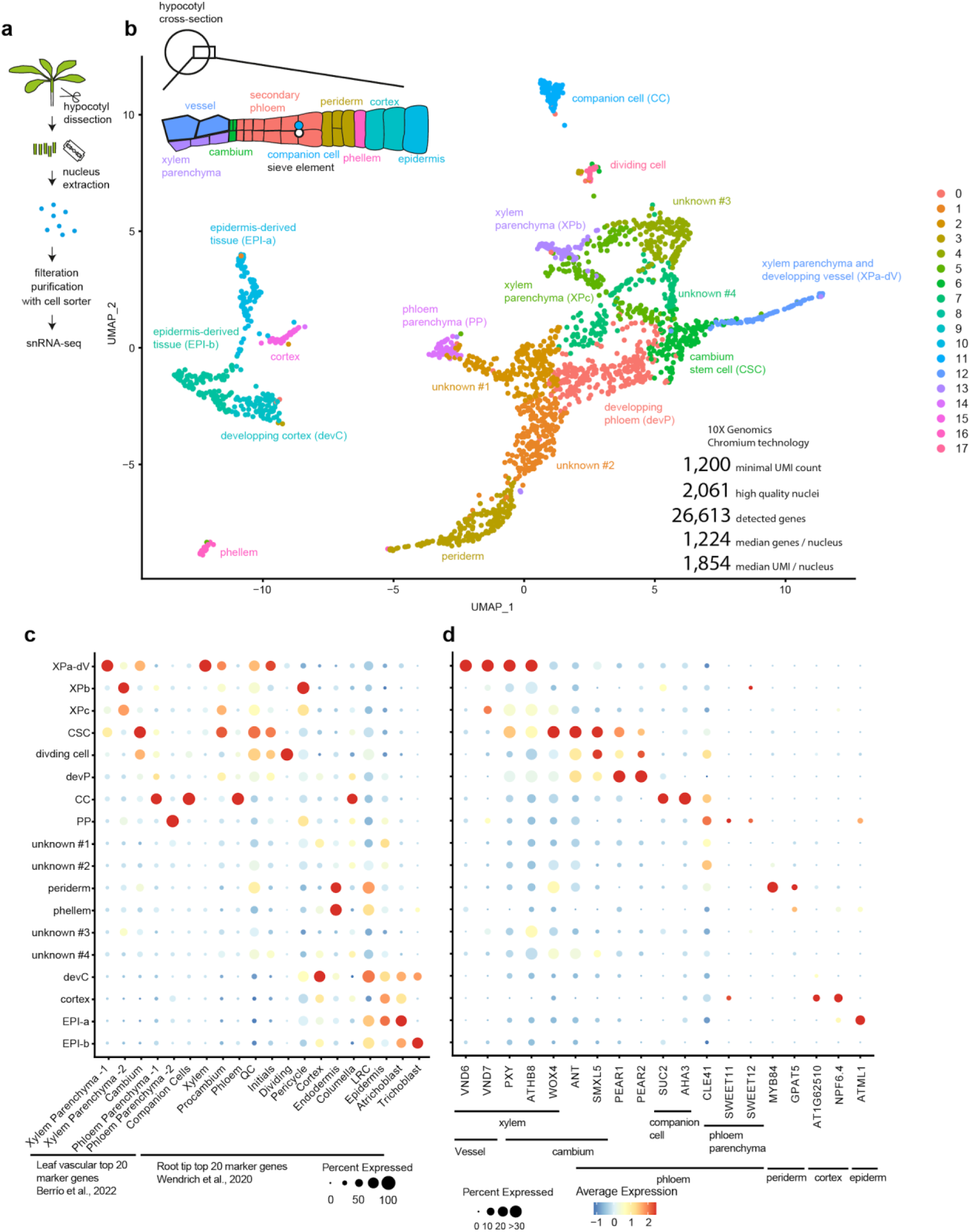
Identification of Arabidopsis hypocotyl cell types using 10× Chromium snRNA-seq. **a**, Sample collection pipeline. **b,** UMAP plot of 10× Chromium snRNA-seq analysis using 2,061 Arabidopsis hypocotyl nuclei organised in 18 clusters obtained through unsupervised clustering analysis. A scheme of major hypocotyl cell types is shown on the top left corner. **c**, **d**, Dot plot showing the expression of tissue-specific genes identified in previous scRNA-seq analyses^12,13^ (**c**) and previously characterised tissue-specific marker genes (**d**) in the identified clusters. The size of the circles represents the percentage of cells with expression (percent expressed), whereas the colour indicates the scaled average expression (average expression).

By unsupervised clustering of the high-quality nuclei, we obtained 18 clusters, as visualised in a ‘uniform manifold approximation and projection’ (UMAP, Fig. 1b, Supplementary Data 2). Based on publicly available cell type-specific marker genes^12,13^, we annotated cluster identities representing all major cell types reported for the hypocotyl (Fig. 1c, d, Supplementary Data 3). These cell types included the cambium (see below) and the periderm, which is a secondary protective tissue characterised by *MYB84*, *GLYCEROL-3-PHOSPHATE SN-2-ACYLTRANSFERASE5* (*GPAT5*), and *WUSCHEL RELATED HOMEOBOX4* (*WOX4*) expression^14^. As an exception, phloem sieve elements were not detected, presumably due to the absence of nuclei in those cells. Importantly, we identified a CSC-associated cluster containing 139 nuclei expressing 151 marker genes including the known CSC markers *WOX4*^6,15,16^, *PHLOEM INTERCALATED WITH XYLEM* (*PXY*)^6,17,18^, *SUPPRESSOR OF MAX2 1-LIKE5* (*SMXL5*)^6,19^, *AINTEGUMENTA* (*ANT*)^4,20^, and *ARABIDOPSIS THALIANA HOMEOBOX8* (*ATHB8*)^21^ (p-value adjusted based on Bonferroni correction (p_adj) < 0.01, Fig. 1d, Supplementary Data 2)^22^. The CSC cluster was surrounded by clusters representing differentiating xylem and phloem cells (Fig. 1b). This was in remarkable contrast to previous analyses of procambium cells, the precursors of cambium cells during apical growth, which usually localise at the end of developmental trajectories^13^. Collectively, our results revealed that CSCs hold a specific transcriptome and reflected the central role of CSCs as the developmental origin of cells driving radial plant growth.

To increase the analytic depth of our CSC characterization, we next applied the plate-based ‘vast transcriptome analysis of single cells by dA-tailing’ (VASA-seq) method^23^ and used fluorescence intensity of the cambium domain reporters *PXY_pro_:H4-GFP* and *SMXL5_pro_:H2B-RFP*^6^ during fluorescence-activated sorting to enrich cambium nuclei (Supplementary Fig. S1c). In total, we processed 2,268 nuclei resulting in 1,559 high quality nuclei containing at least 1,200 unique fragment identifier (UFI) counts/nucleus (Fig. 2a, Supplementary Data 1). Overall, we detected transcripts of 33,195 genes including the two reporter transgenes, covering more than 85 % of the annotated genes in Arabidopsis. On the median level, our VASA-seq approach detected 2,688 genes/nucleus and 5,210 UFI/nucleus, which matched previous high quality datasets for Arabidopsis nuclei^24^.

**Fig. 2:**
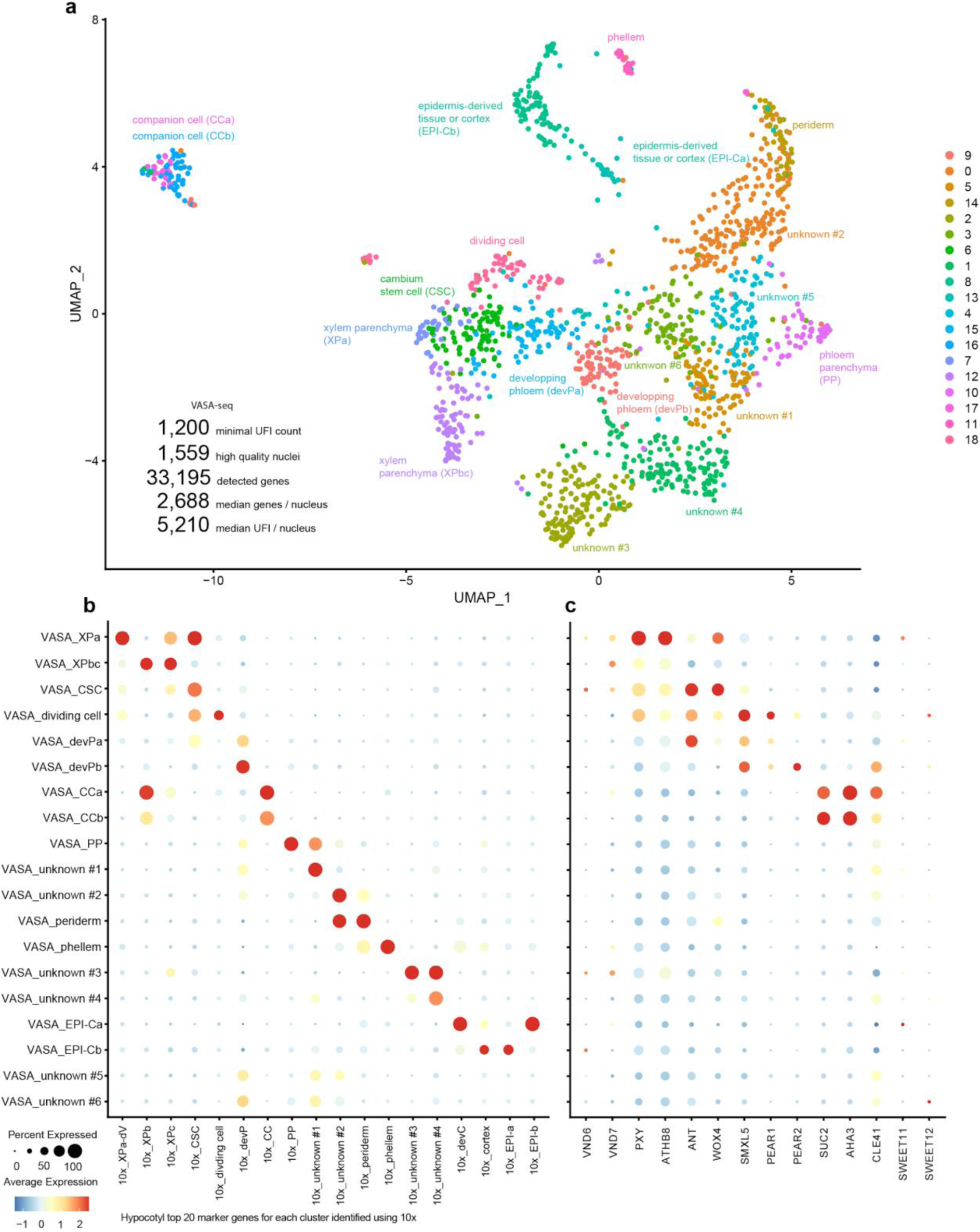
Identification of hypocotyl cell types using VASA-seq. **a**, UMAP plot of VASA-seq analysis using 1,559 hypocotyl nuclei organised in 19 clusters obtained through unsupervised clustering. **b–c**, Dot plots showing the expression of tissue-specific genes identified by 10× Chromium analysis (Supplementary Data 1, 2) (**b**) and previously characterised tissue-specific marker genes (**c**), validating the annotation of cluster identities. The size of circles represents the percentage of cells with expression (percent expressed), whereas the colour indicates the scaled average expression (average expression).

Unsupervised clustering of the VASA-seq processed nuclei resulted in 19 clusters which we associated with distinct cell states using marker genes identified during our droplet-based approach and previously characterised genes (Fig. 2b, c). Again, most hypocotyl cell types were represented by a cluster in the VASA-seq dataset with a lower relative size of ‘periderm’ or ‘developing vessel’ clusters than obtained by our droplet-based approach presumably due to the depletion of those nuclei during the cambium-centred nucleus selection (Supplementary Data 2). Moreover, CSC marker genes were expressed this time in two clusters which represented CSCs (‘VASA_CSC’ cluster, 94 nuclei) and dividing CSCs (‘VASA_dividing’ cluster, 66 nuclei) (Fig. 2b, c) based on transcript abundance of genes previously associated with dividing root cells^13^. Supporting the conclusion that these were CSC nuclei, both clusters showed high GFP (*PXY_pro_:H4-GFP*) and RFP (*SMXL5_pro_:H2B-RFP*) fluorescence intensities^6^ (Supplementary Fig. S2a). We furthermore identified 226 and 405 marker genes for each CSC cluster, respectively (p_adj < 0.01, Supplementary Data 2) demonstrating that the VASA-seq analysis substantially increased analytical depth. Moreover, 119 of 151 (72 %) CSC marker genes identified by our droplet-based approach were included in the groups of CSC and/or dividing CSC marker genes (Supplementary Fig. S2b), suggesting that results from both approaches were robust.

### SL signalling activity within radial growth

Demonstrating their relevance, our cell-resolved transcriptome data recapitulated xylem-associated auxin signalling and phloem-associated cytokinin signalling^25^ when we monitored transcript abundance of respective hormone-responsive genes^26^ within identified clusters (Supplementary Fig. S3). Interestingly, when we mapped the activity of 94 genes induced by the synthetic SL analogue GR24^4DO, 27^, we found that the expression of these genes was particularly low in dividing CSCs and in a subset of xylem cells, presumably differentiating vessel elements (‘VASA_XPa’, Fig. 3a, b). This finding suggested differential SL-signalling activity during radial growth-associated cell fate transitions. Indeed, when analysing SL signalling levels by the genetically encoded SL signalling sensor Strigo-D2^28^, we found that SL-signalling activity is lower in cambium cells and in developing vessel elements compared to phloem and xylem parenchyma cells (Fig. 3c–e).

**Fig. 3:**
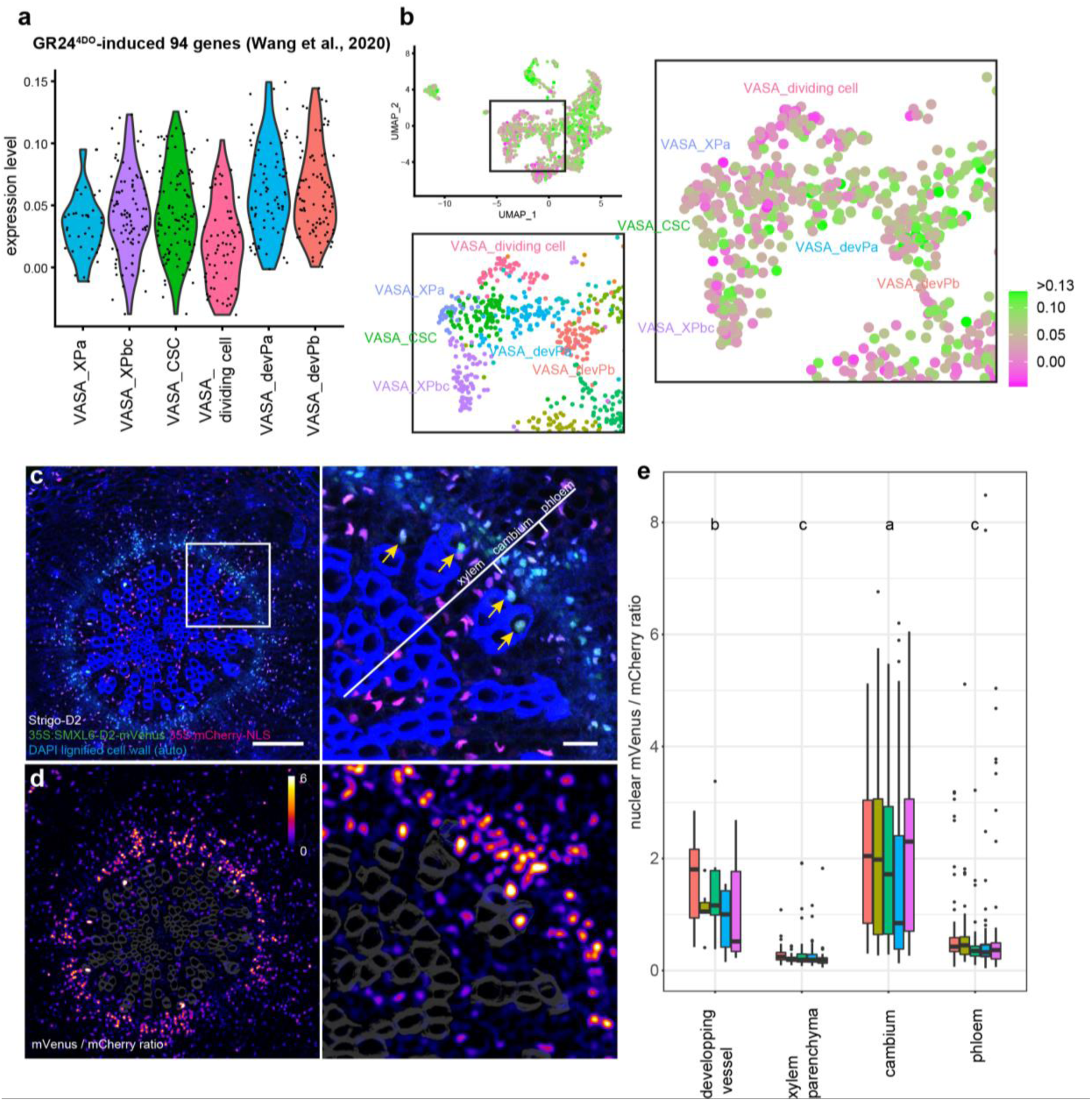
Strigolactone (SL) signalling in the radially expanding hypocotyl. **a**, **b**, Violin plot (**a**) and UMAP visualisation (**b**) of expression profiles of 94 GR24^4DO^-induced genes^27^ in the cambium-related cell clusters identified by VASA-seq. Statistical groups determined by the Steel-Dwass test for multiple comparisons are indicated by letters (*p* < 0.05). Gene lists used in this analysis can be found at Supplementary Data 3. A close-up of the UMAP originated from the VASA-seq cluster identification (Fig. 2) is shown as a reference. **c,** Maximum intensity projection of confocal microscopy images of 5-week-old hypocotyl cross-sections expressing Strigo-D2 (*35S_pro_:SMXL6-D2-mVenus_35S_pro_:mCherry-NLS*) as an indicator of SL signalling. mVenus and mCherry signals are shown in green and magenta, respectively. Hoechst33342 was used to stain nuclei shown in blue together with the autofluorescence of lignified cell walls in vessel elements. Yellow arrowheads indicate the nuclei of developing vessel elements. Size bars represent 100 µm and 20 µm on the left and right, respectively. **d**, Ratiometric images of mVenus / mCherry signals after image processing (see Methods). The colour bar indicates the scale of ratio values from 0 to 6. Higher ratio indicates lower SL signalling levels. **e**, Box plot showing nuclear fluorescence intensity ratio of mVenus and mCherry in different vascular tissues. Data from one individual plant are shown in the same colour (n=5). The total number of nuclei analysed for each tissue type of five plants was 42 (developing vessels), 389 (xylem parenchyma), 903 (cambium), and 411 (phloem). Statistical groups are indicated by letters and assessed by a one-way ANOVA with post-hoc Tukey-HSD (95 % CI) on the mean ratio value of each tissue type in each individual plant.

SLs are a class of carotenoid-derived phytohormones, modulating various processes including shoot branching and drought resistance^29,30^. SL molecules are perceived by the α/β hydrolase DWARF14 (D14) inducing interaction with the F-box protein MORE AXILLARY GROWTH2 (MAX2) and thereby promoting mainly degradation of the transcriptional regulators SMXL6, SMXL7, and SMXL8^27,31,32^. Using fluorescent promoter reporters, we found that the genes encoding these signalling components were widely active in the hypocotyl with a particular focus on the xylem (Fig. 4): The activity of the *MAX2* reporter was detected in developing xylem vessels, xylem parenchyma cells, cambium cells and phloem parenchyma cells, whereas *D14* reporter activity was only detected in xylem parenchyma cells and phloem parenchyma cells. *SMXL6*, *SMXL7*, and *SMXL8* reporter activities were all detected in xylem parenchyma cells, with the *SMXL7* reporter also being detected in developing vessels (Fig. 4). Taken together, our analysis associated SL signalling components with differentiating and differentiated hypocotyl tissues and were in line with a role of the pathway during the formation of vascular cell types.

**Fig. 4:**
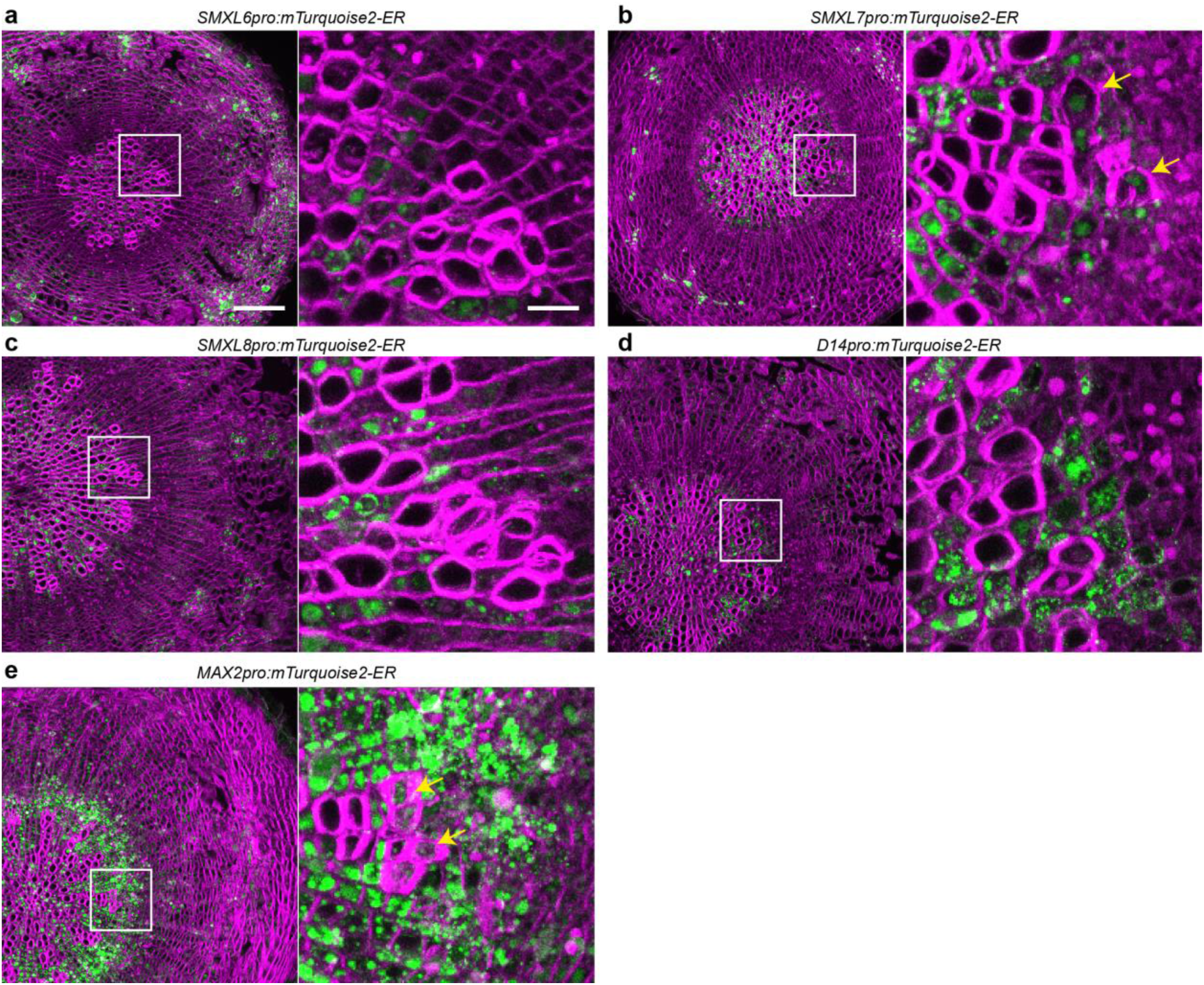
Activity of transcriptional reporters for genes involved in SL signalling in hypocotyl sections. **a–e**, Maximum intensity projection of confocal images obtained from hypocotyl cross-sections of 4-week-old plants carrying *SMXL6_pro_:mTurquoise2-ER* (**a**), *SMXL7_pro_:mTurquoise2-ER* (**b**), *SMXL8_pro_:mTurquoise2-ER* (**c**), *D14_pro_:mTurquoise2-ER* (**d**), or *MAX2_pro_:mTurquoise2-ER* (**e**) transgenes, respectively. mTurquoise2 signals are shown in green. Cell walls were stained by Direct Red 23 and are shown in magenta. Magnified images of white squared regions are shown on the right of each figure. Yellow arrows indicate developing vessel elements. Scale bars represent 100 μm on the left images, and 20 μm in the magnified images on the right.

### SL-signalling suppresses vessel formation

To reveal a possible function of the SL-signalling pathway in vascular tissue formation, we next compared hypocotyls from *d14* mutants and wild type by again applying droplet-based snRNA-seq. After clustering and cluster annotation based on our initially identified genes marking the different hypocotyl cell types (Fig. 5a, c, Supplementary Data 1), we found that cell composition and transcriptional states were highly similar between the two genotypes (Fig, 5b, d). However, as the most prominent difference, we detected a 3-fold increase in the abundance of nuclei from developing vessel elements (WTd14_devV) in *d14* mutants (Fig. 5e). Those were identified based on the expression of the *VASCULAR-RELATED NAC-DOMAIN6* (*VND6*), *VND7^33,34^* and *IRREGULAR XYLEM3* (*IRX3)^35^* and other xylem marker genes (Fig. 5f, Supplementary Fig. S4). In line with an increased number of vessel elements, transcripts of *VND6*, *VND7* and *IRX3* were more abundant in *d14* mutants (Fig. 5g). Indeed, histological analyses revealed a higher density and increased size of vessel elements in both the SL signalling mutants *d14* and *max2* (Fig. 6). This alteration was not observed in *BRANCHED1* (*BRC1*)-defective plants (Fig. 7a–c) which, similarly as *d14* and *max2* mutants, show increased branching^36^ arguing against the possibility that increased shoot formation causes enhanced vessel formation in *d14* and *max2* as a secondary effect. In contrast to the two SL signalling mutants, the number and size of vessel elements was decreased in *smxl6;smxl7;smxl8* (*smxl6;7;8*) and *d14*;*smxl6;7;8* mutants (Fig. 6) demonstrating that increased vessel formation in *d14* depends on the proteolytic targets of the SL signalling pathway. This conclusion was supported when determining the vessel area to xylem area ratio in *max2;smxl6;7;8* quadruple mutants which was, in contrast to an increased density in *max2* single mutants, comparable to wild type plants (Supplementary Fig. S5). To this effect, mostly *SMXL7* and *SMXL8* contributed in a redundant fashion, as no other *max2;smxl* double or triple mutant combination than *max2;smxl7;8* showed a mild reduction in vessel density in comparison to *max2* (Supplementary Fig. S5). Further supporting a positive role of the proteolytic targets of SL-signalling in vessel formation, expression of an SL-resistant version of SMXL7 (SMXL7^d53^)^37^ under the control of the *SMXL7* promoter promoted cambium-associated vessel formation (Fig. 7g–i). Indicating a vascular-associated role of SL signalling, *d14* mutants did not show an increased vessel phenotype when they carried a transgene expressing *D14* under the control of the cambium-and early xylem-specific *WOX4* promoter (Fig. 7d–i). Together with the observation that application of GR24^4DO^ decreased vessel formation (Fig. 8), we concluded that SL-signalling negatively regulates vessel formation during radial plant growth in cambium-related cells and in an *SMXL6;7;8*-dependent fashion.

**Fig. 5:**
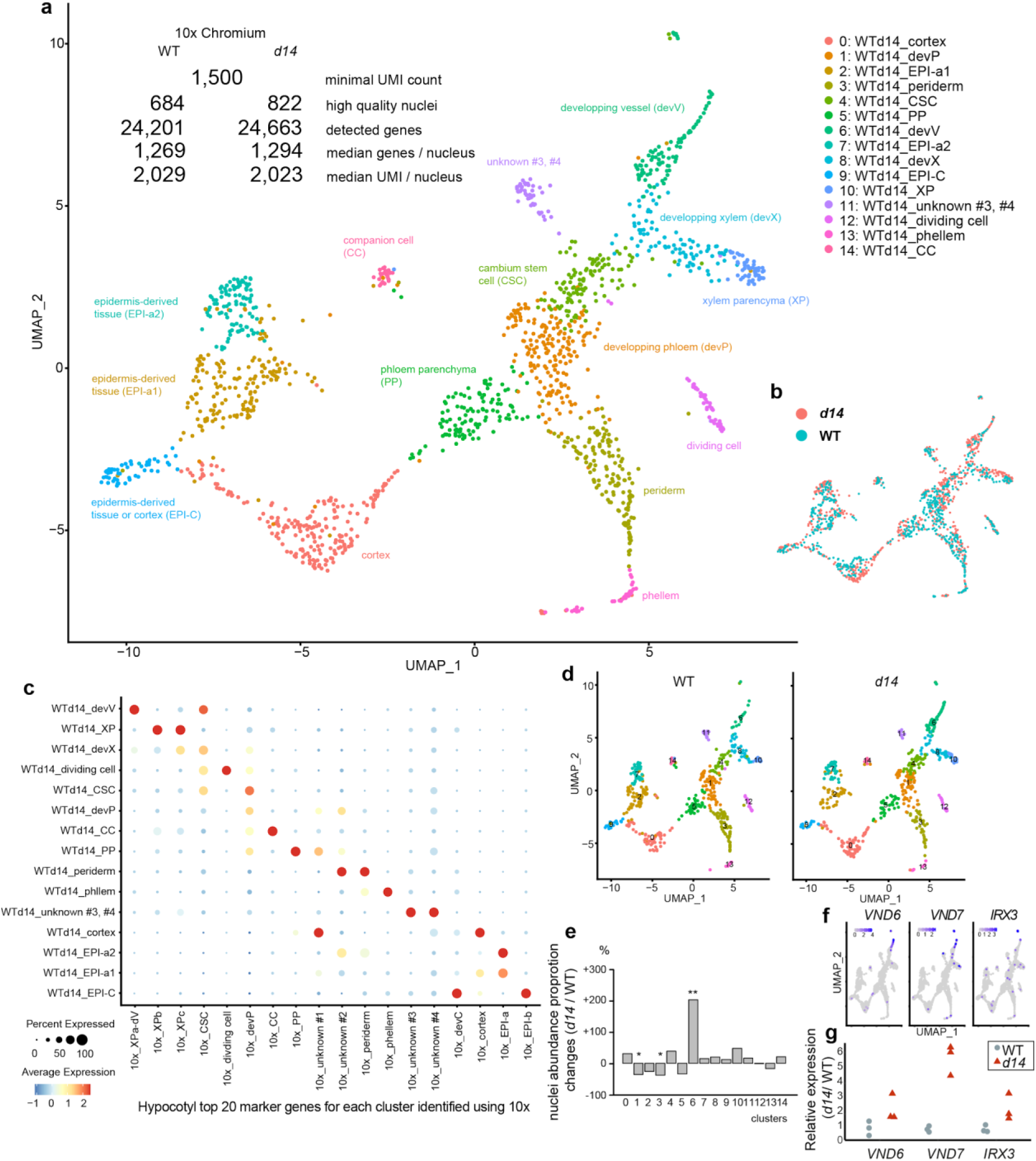
Identification of hypocotyl cell types applying 10× Chromium on wild type and *d14* mutants. **a**, UMAP plot of snRNA-seq analysis using in total 1,506 nuclei collected from wild type and *d14* mutants organised in 15 clusters obtained through unsupervised clustering. **b**, UMAP plot colour-coded by genotype. **c**, Dot plot showing the expression of tissue-specific genes identified by 10× Chromium analysis (Supplementary Data 2, Supplementary Data 3). The size of the circles represents the percentage of cells with expression (percent expressed), whereas the colour indicates the scaled average expression (average expression). **d**, UMAP plot of 10× Chromium snRNA-seq analysis using 500 hypocotyl nuclei from (**a**). 500 nuclei were randomly resampled from 684 (WT) and 822 (*d14*) nuclei, respectively. **e**, Changes in relative nuclei abundance for each cluster in *d14* mutants compared to wild type. Fisher’s exact test was used to determine statistical differences in each cluster between genotypes. * *p*<0.05, ** *p*<0.01. *p* = 1.0e-03 (cluster 1:WTd14_devP), *p* = 3.1e-03 (cluster 3:WTd14_periderm), *p* = 2.9e-07 (cluster 6:WTd14_devV). **f**, Expression of the developing vessel marker genes *VND6*, *VND7* and *IRX3* in the UMAP plot shown in (**a**). **g**, Relative transcript abundance of *VND6*, *VND7* and *IRX3* in *d14* mutants compared to wild type, normalised to the *EF-1a* reference gene in qRT-PCR experiments. n = 3 biological replicates.

**Fig. 6:**
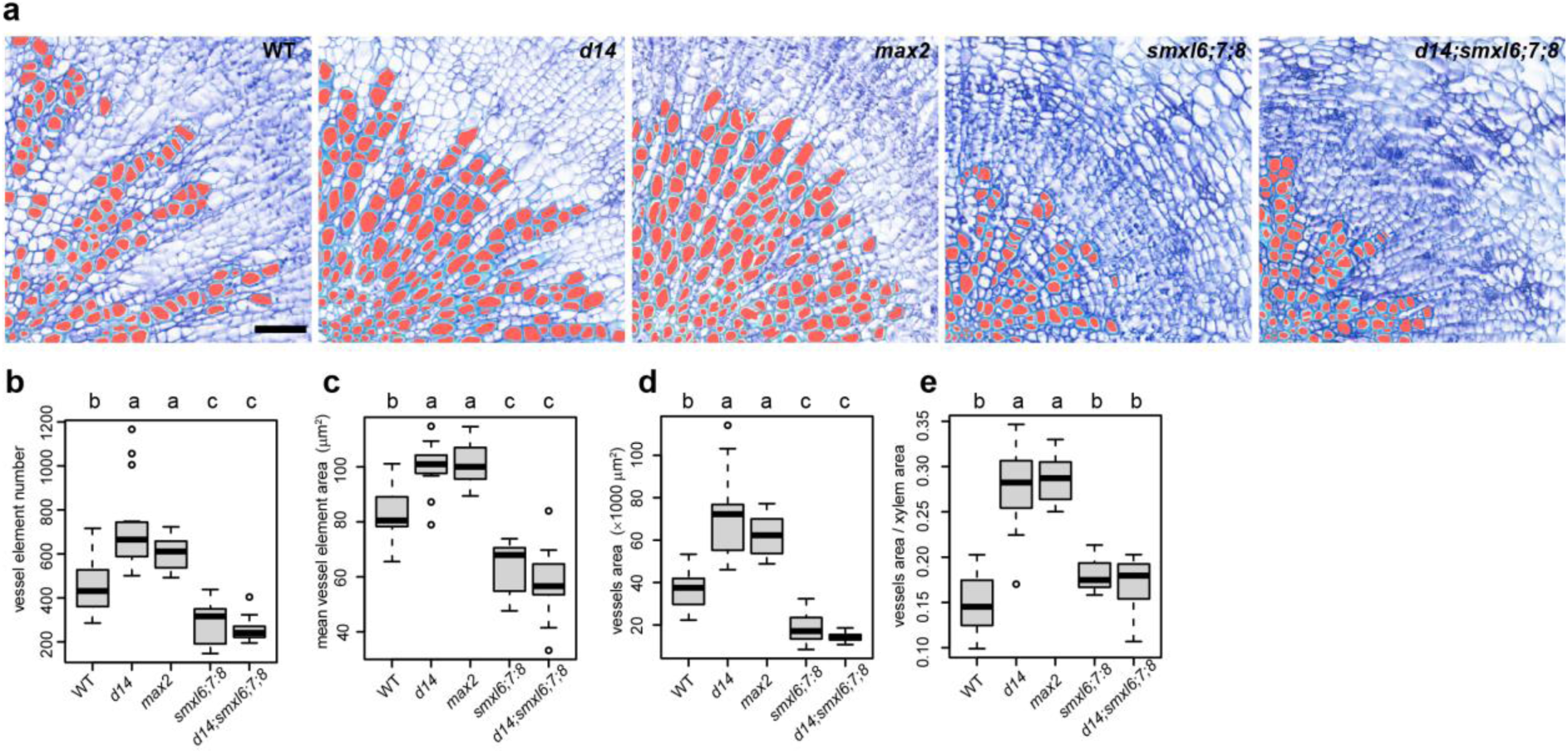
Effect of SL signalling on vessel development. **a**, Toluidine-blue stained hypocotyl cross-sections from 5-week-old wild type, *d14*, *max2*, *smxl6;7;8* and *d14;smxl6;7;8* plants. Vessels were automatically identified using ImageJ and subsequent manual correction (marked in red). Scale bar represents 50 μm. **b**–e, Quantification of the vessel element numbers per section (**b**), the average area of individual vessel elements (**c**), the total vessel area per section (**d**) and the ratio between the vessel element area and the total xylem area (**e**) in different genotypes. n=10-15 plants for each genotype. Statistical groups are indicated by letters and determined by a one-way ANOVA with post-hoc Tukey-HSD (95% CI).

**Fig. 7:**
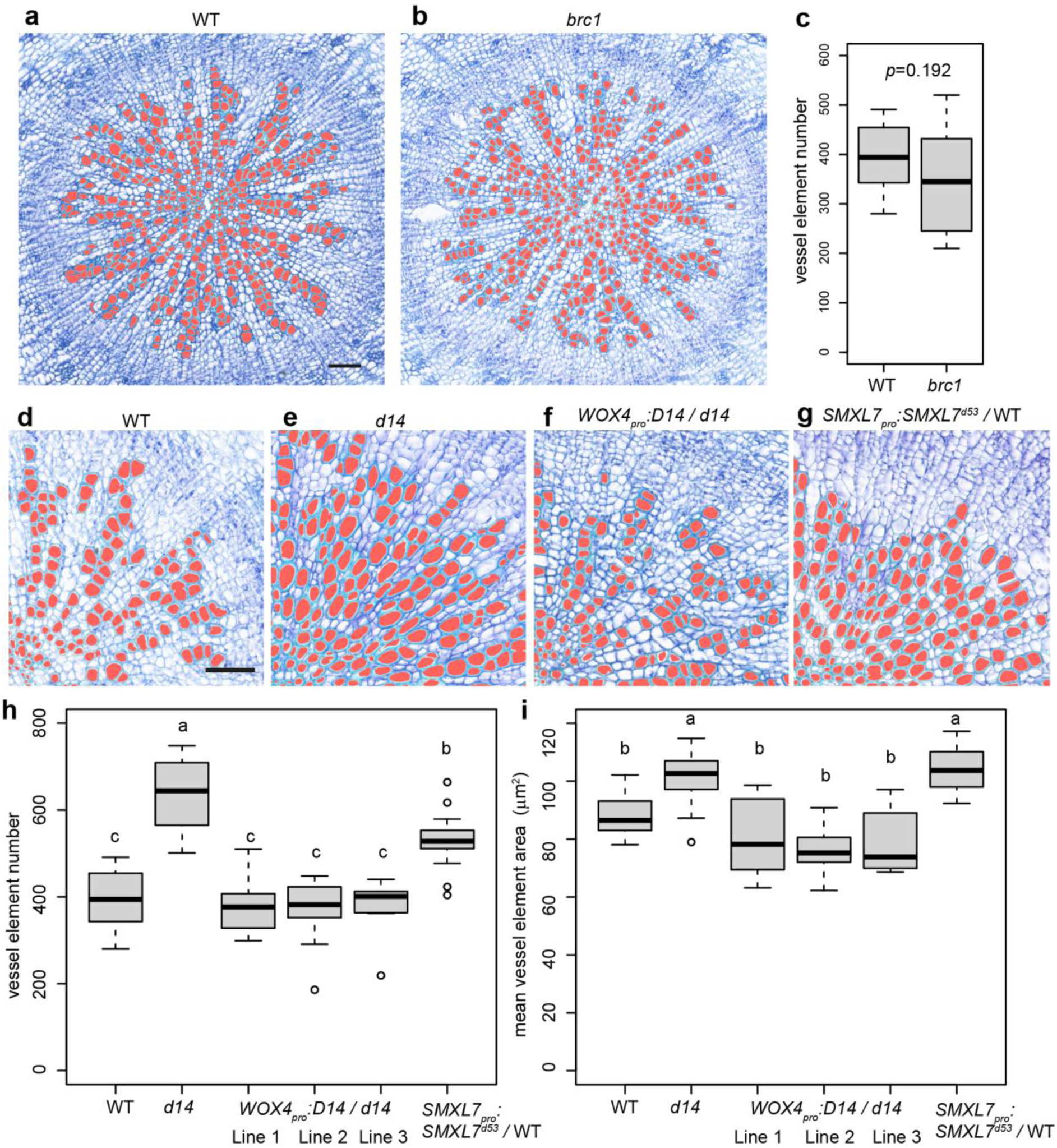
Analysis of *brc1*, *SMXL7_pro_:SMXL7^d53^-VENUS*, and *WOX4_pro_:D14;d14*plants. **a**, **b**, Toluidine blue-stained hypocotyl cross-sections from 5 weeks-old wild type (**a**) and *brc1* (**b**) plants. **c**, Quantification of vessel elements per section comparing wild type and *brc1* plants. n=11 (wild type) and 10 (*brc1*). *p* value was determined by the Welch’s t-test. **d**–**i**, Toluidine blue-stained hypocotyl cross-sections from 5 week-old wild type (**d, g**) and *d14* (**e, f**) plants carrying a *WOX4_pro_:D14* (**f**) or *SMXL7_pro_:SMXL7^d53^-VENUS* transgene (**g**). Vessel elements were automatically identified using ImageJ and subsequent manual correction (marked in red). **h**, **i**, Quantification of vessel element number per section (**h**) and mean vessel element area size per section (**i**) comparing wild type, *d14*, three independent lines carrying *WOX4_pro_:D14* transgenes in the *d14* background and lines carrying a *SMXL7_pro_:SMXL7^d53^-VENUS* transgene (n=7-13 plants for each genotype). Statistical groups are indicated by letters and were determined by a one-way ANOVA with post-hoc Tukey-HSD (95 % CI). Scale bar represents 50 μm.

**Fig. 8:**
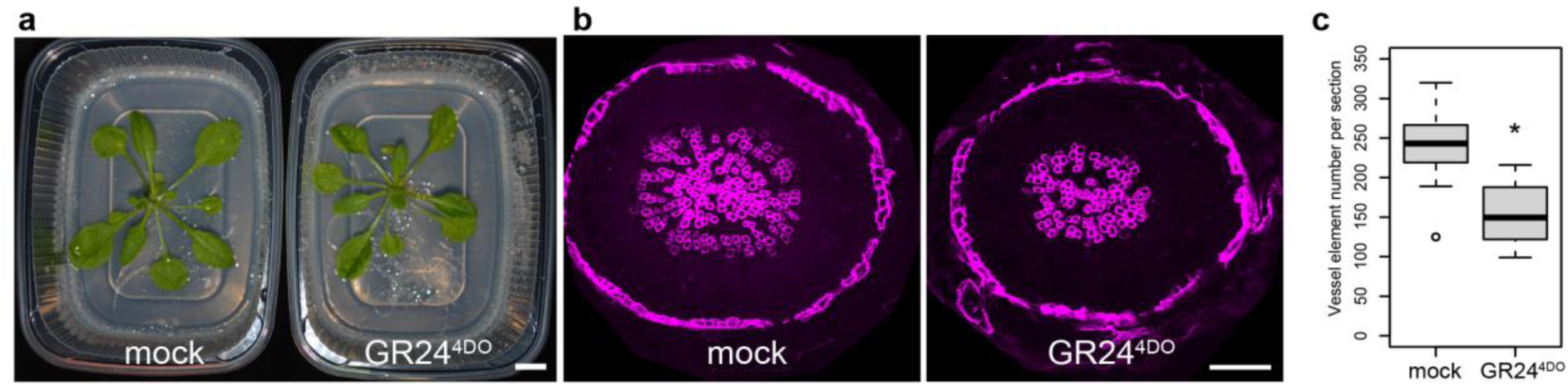
Histological analysis of the GR24^4DO^ effect on vessel formation. **a**, Appearance of 4-week-old plants after application of 10 μM acetone (left, mock) or GR24^4DO^ (right). Scale bar represents 1 cm. **b**, Hypocotyl cross-sections of 4-week-old plants after application of 10 μM acetone (left) and GR24^4DO^ (right). The cross sections were excited using 405 nm laser light, and the auto-fluorescence from the lignified vessel elements was captured and displayed in magenta. The peripheral periderm auto-fluoresces as well due to suberin deposition. Scale bar represents 100 μm. **c**, Quantification of vessel elements per section comparing plants treated with 10 μM acetone and GR24^4DO^. n=15 (mock), n=14 (GR24^4DO^) plants. Asterisk indicates *p*<0.01 determined by the Welch’s t-test (*p*=2.5e-05).

### Vessel development influences transpiration

Relative water content is reduced in *d14* and *max2* mutants^38–40^ and increased in *smxl6,7,8* triple mutants when grown in water-deprived pots^41^, suggesting that SL signalling optimises water usage. Confirming this role, water usage was increased in *d14* and *max2* mutants and reduced in *smxl6;7;8* mutants (Fig. 9a). Abscisic acid (ABA)-induced closure of stomata, the main transpiration site in plants, is slower in SL signalling deficient mutants and proposed as one cause for their reduced drought resistance^38–40^. Indeed, stomata conductance was elevated in *d14* and *max2* mutants and lower in *smxl6;7;8* mutants compared to wild type plants under both well-watered and water deficiency conditions (Fig. 9b), demonstrating that transpiration is reduced by SL signalling. Again indicating that this effect depended on SL signalling in vascular tissues, *d14* mutants carrying a *WOX4_pro_:D14* transgene (see above) showed normal transpiration rates in leaves (Supplementary Fig. S6a, c). In accordance with previous reports^39,40^, stomata density was not significantly changed in *d14* and *max2* mutants and only slightly enhanced in *smxl6;7;8* mutants (Supplementary Fig. S6d, e) arguing against altered stomata density as a reason for altered transpiration in SL-related mutants.

**Fig. 9:**
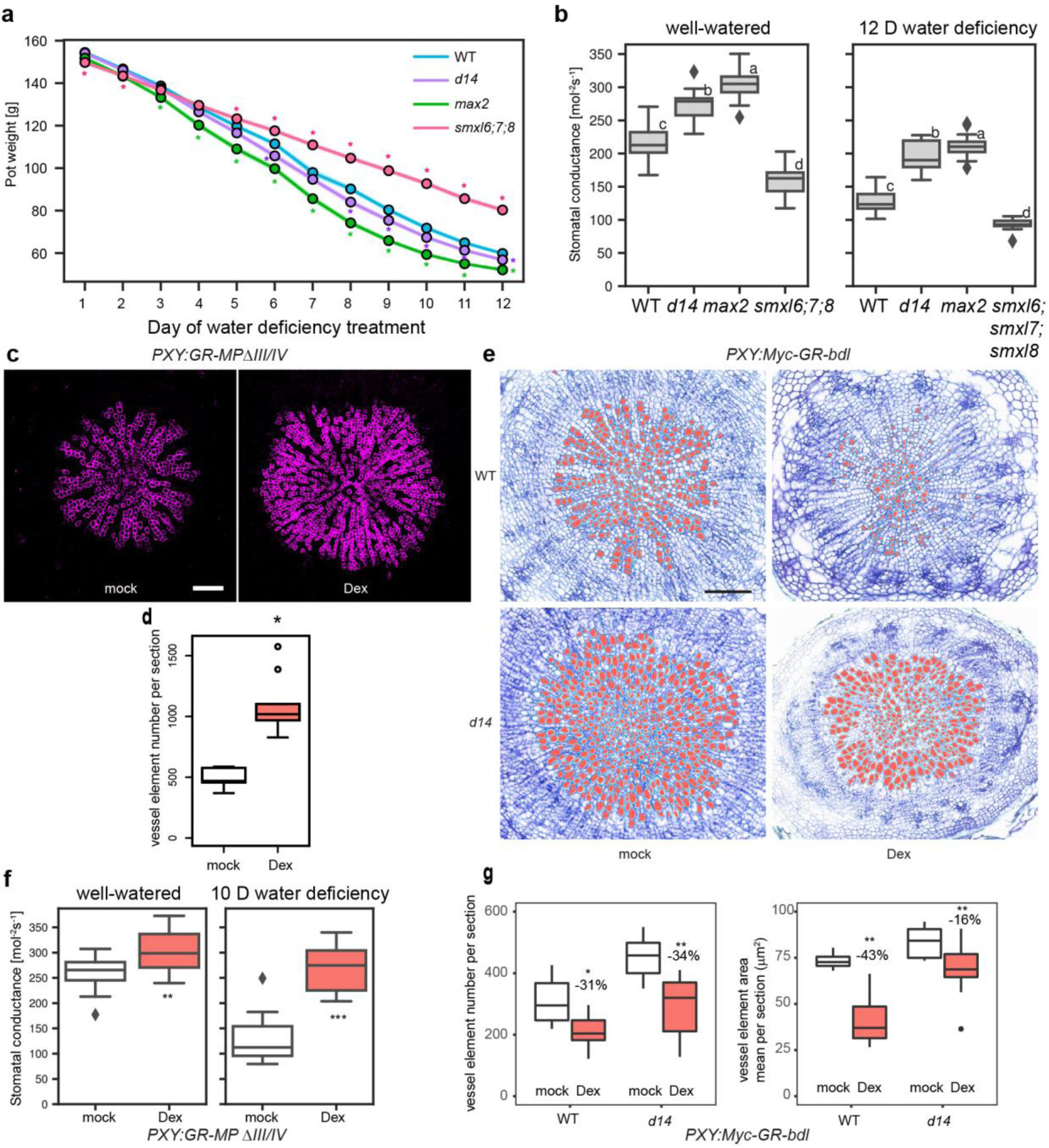
Effect of altered vessel formation on water usage and stomatal conductance. **a**, Pot weight changes when growing wild type, *d14*, *max2* and *smxl6;7;8* plants during water deficiency treatments. Measurements were analysed by t-test at each timepoint comparing mutant genotypes to wild type. Asterisks indicate significance (p < 0.01, Supplementary Data 6). n=109 (wild type), n=111 (*d14*), 109 (*max2*), n=108 (*smxl6;7;8*). Results from three independent experiments are included. **b**, Quantification and comparison of stomatal conductance found in wild type, *d14*, *max2* and *smxl6;7;8* plants under well-watered conditions and 12 days (12 D) after initiating water deficiency treatments (n=17-19). Conductance was measured on three leaves per plant. Plot shows average conductance of the three leaves per plant. **c**, Hypocotyl cross-sections of five week-old plants carrying a *PXY_pro_:GR-MPΔIII/IV* transgene treated with mock or Dex solutions for the last two weeks. Vessel elements are visualised through their autofluorescence when excited by the UV laser (shown in magenta). Scale bar represents 50 μm. **d**, Vessel element numbers per section comparing Dex and mock-treated *PXY_pro_:GR-MPΔIII/IV* plants as shown in c (n=9 plants). The asterisk indicates p<0.01 when the Welch’s t-test was applied (p=3.2e-05). **e**, Hypocotyl cross-sections from 31 day-old wild type and *d14* plants carrying the *PXY_pro_:Myc-GR-bdl* transgene. Dex was applied from three weeks onwards for 10 days. Semi-automatically detected vessel elements are marked in red. Scale bar represents 100 μm. **f**, Quantification and comparison of stomatal conductance in *PXY_pro_:GR-MPΔIII/IV* plants treated with mock or Dex solution under well-watered or water-deficiency (10 days) conditions. Stomatal conductance was measured on 3 leaves per plant with an SC1 Leave prometer. n=15-18 plants for each genotype. The asterisk indicates significance when the student’s t-test was applied, ** p < 0.01, *** p < 0.001, p=2.7e-03 (well-watered), p=4.94e-09 (10 days water deficiency). **g**, Vessel element numbers per section (left) and the mean of vessel element area per section (right) in response to Dex treatment comparing wild type and *d14* mutants as shown in e (n=8-15 plants). * p<0.05, ** p<0.01 in the post-hoc Bonferroni test after two-way ANOVA for the effect of treatment for each genotype. *p*=0.022 (number, WT), *p*=1.4e-05 (number, *d14*), *p*=1.5e-06 (area, WT), *p*=0.0033 (area, *d14*). Two-way ANOVA analysis indicates that there is an effect of the treatment on vessel element number which is independent from genotypes (F=1.334, df=1, p=0.254), as well as on vessel element area which is dependent on genotypes (F=6.578, df=1, p=0.014).

Based on these findings, we rationalised that SL signalling restricts water usage in plants by modulating vessel formation and, consequently, water transport capacity. To see to which extent the modulation of vessel formation influences transpiration, we perturbed vessel formation by altering auxin signalling. AUXIN RESPONSE FACTOR (ARF) transcription factor-mediated auxin signalling promotes cambium-derived vessel formation^4^ with MONOPTEROS (MP, also known as ARF5), playing an important role in this context^4,42^. In accordance with previous findings^4^, induction of a dexamethasone (dex)-inducible and auxin-insensitive version of MP, a glucocorticoid receptor (GR)-MPΔIII/IV fusion protein^43^, expressed under the control of the cambium-associated *PXY* promoter^42^ resulted in a significant increase of vessel formation (Fig. 9c, d). In contrast, when auxin signalling levels were suppressed in *PXY*-positive cells by inducing the GR-linked auxin signalling repressor bodenlos (GR-bdl)^42,44,45^, vessel formation was reduced (Fig. 9e, g). This effect was weaker in *d14* mutants suggesting that SL signalling is required for auxin-dependent vessel formation. Supporting an impact of the number of vessel elements on water usage, we detected elevated stomata conductance in dex-induced *PXY_pro_:GR-MPΔIII/IV* plants compared to mock-treated plants in both well-watered and water deficiency condition (Fig. 9f). This effect was observed although stomata density was not significantly affected upon induction (Supplementary Fig. S6f, g). These results supported our hypothesis that vessel abundance directly influences stomata conductance.

Taken together, by starting off with a cell-resolved transcriptome analysis of radially growing Arabidopsis organs, we revealed that SL signalling modifies the number and size of vessel elements produced during cambium-dependent radial growth. The possibility to modulate vessel anatomy in response to environmental cues like drought^46^, can be assumed to be an important fitness trait and has, so far, been described for being mediated by abscisic acid (ABA) signalling during longitudinal growth^47^. We did, however, neither observe any alteration of primary vessel element formation in primary *d14* roots (Fig. 10a), nor an increase of cambium-derived vessel formation in *aba deficient 2* (*aba2*) mutants (Fig. 10d, f) impaired in ABA biosynthesis^48^ and showing higher stomata conductance^49^. Moreover, the formation of cambium-derived vessel elements was not altered in mutants displaying an increased stomata density and a resulting increase in stomata conductance^50,51^ (Fig. 10e, f). These findings suggest that ABA-and SL-signalling pathways fulfil distinct functions with regard to their effect on vessel formation during primary and secondary development, respectively. Specifically, we propose that increased stomata conductance in SL-related mutants is a result of enhanced vessel element formation and an associated increase of water transport along the vascular system. Interestingly, SL biosynthesis is enhanced in rice roots in drought conditions^52^ indicating that the pathway has the potential to lead to structural changes when plants experience growth, to possibly anticipate harsh conditions in the future. Depending on distinct ecological or agricultural niches plants have adapted to, it may be advantageous to establish a weaker or stronger impact of SL signalling on vessel element formation to cope with distinct environmental regimes.

**Fig. 10:**
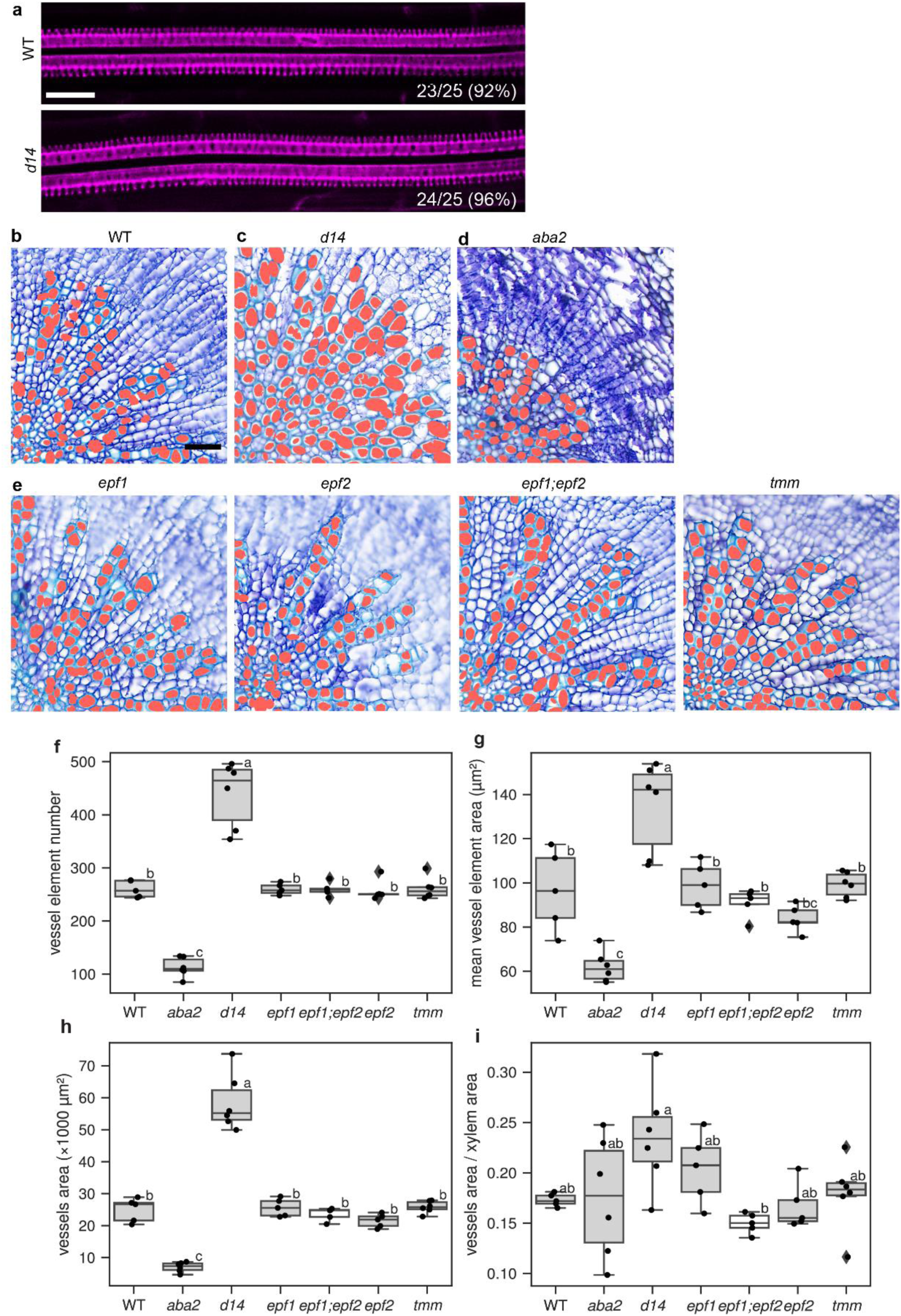
Primary vessel development in *d14* mutants and cambium-derived vessel formation in ABA signalling-and stomata-related mutants. **a**, Morphology of xylem strand in wild type and *d14* roots five days after germination. Xylem strands were visualised by Basic Fuchsin staining. The proportion indicates the frequency of plants observed with two metaxylem and two protoxylem strands. The scale bar represents 20 μm. **b**–**e**, Toluidine blue-stained hypocotyl cross-sections from five week-old wild type (WT, **b**) and *d14* (**c**), ABA biosynthesis mutant *aba2-11* (**d**) and stomata-related mutants (**e**) including *epidermal patterning factor 1* (*epf1*), *epf2*, *epf1;epf2*, and *too many mouths* (*tmm*). Scale bar represents 50 μm. **f-i**, Quantification of the vessel element numbers per section (**f**), the average area of individual vessel elements (**g**), the total vessel area per section (**h**) and the ratio between the vessel element area and the total xylem area (**i**) in each mutant shown in (**b**– **e**). n=5-6 plants for each genotype. Statistical groups are indicated by letters and were determined by a one-way ANOVA with post-hoc Tukey-HSD (95 % CI).

## Acknowledgements

10× Chromium operation, library preparation and next-generation sequencing were carried out by David Ibberson (The CellNetworks Deep Sequencing Core Facility, Heidelberg University, Germany). Nucleus sorting was carried out by Monika Langlotz (Flow Cytometry and FACS Core Facility, ZMBH, Heidelberg University, Germany). VASA-seq processing was carried out by Single Cell Discoveries B.V. (Utrecht, The Netherlands). We thank David Nelson (University of California Riverside, USA), Christopher Grefen (University of Bochum, Germany), Mikael Brosché (University of Helsinki, Finland), Ottoline Leyser (University of Cambridge, UK) for providing plant material, Virginie Jouannet (Heidelberg University, Germany) for generating plant material and Jan Lohmann (Heidelberg University, Germany) for sharing equipment. This work was supported by a fellowship of the Chinese Scholarship Council (CSC) and a Peterson scholarship (http://www.peterson-elites.sdu.edu.cn/jxjjj.htm) to J.Z., an overseas research fellowship of the Japan Society for the Promotion of Science [JSPS Overseas 201960008] and a JST PRESTO grant [JPMJPR2046] to D.S.. Ki.K. was supported by a doctoral fellowship of the Landesgraduiertenförderung of the state Baden-Württemberg. Work in T.G.’s group was furthermore supported by the SFB 873 and SFB 1101 of the Deutsche Forschungsgemeinschaft (DFG) and the ERC consolidator grant PLANTSTEMS. Work in the Ke.K. group was supported by the DFG project 438774542.

## Author contributions

J.Z., D.S., Ki.K. and T.G. designed the study. J.Z., D.S., Ki.K., T.B., and L.L. performed experiments. J.Z., D.S., Ki.K. and C.S. analysed the data. X.X. and Ke.K. provided technical support on single nucleus RNA-seq analysis. J.Z., D.S. and T.G. wrote the manuscript with input from all the authors.

## Data availability

The raw sequencing data of the single nucleus RNA-seq are deposited at NCBI’s Gene Expression Omnibus and are accessible through GEO Series accession number GSE224928 at https://www.ncbi.nlm.nih.gov/geo/. The authors declare that all other data supporting the findings of this study are mentioned in the main text or the supplementary materials and are available upon request.

## Code availability

General codes were used for analyses and specific parameters were provided in the Methods section. Full codes are available upon request.

## Additional information

Supplementary Information is available for this paper.

Correspondence and requests for materials should be addressed to Dongbo Shi or Thomas Greb.

## Supplementary Figures and Legends

**Supplementary Fig. S1:**
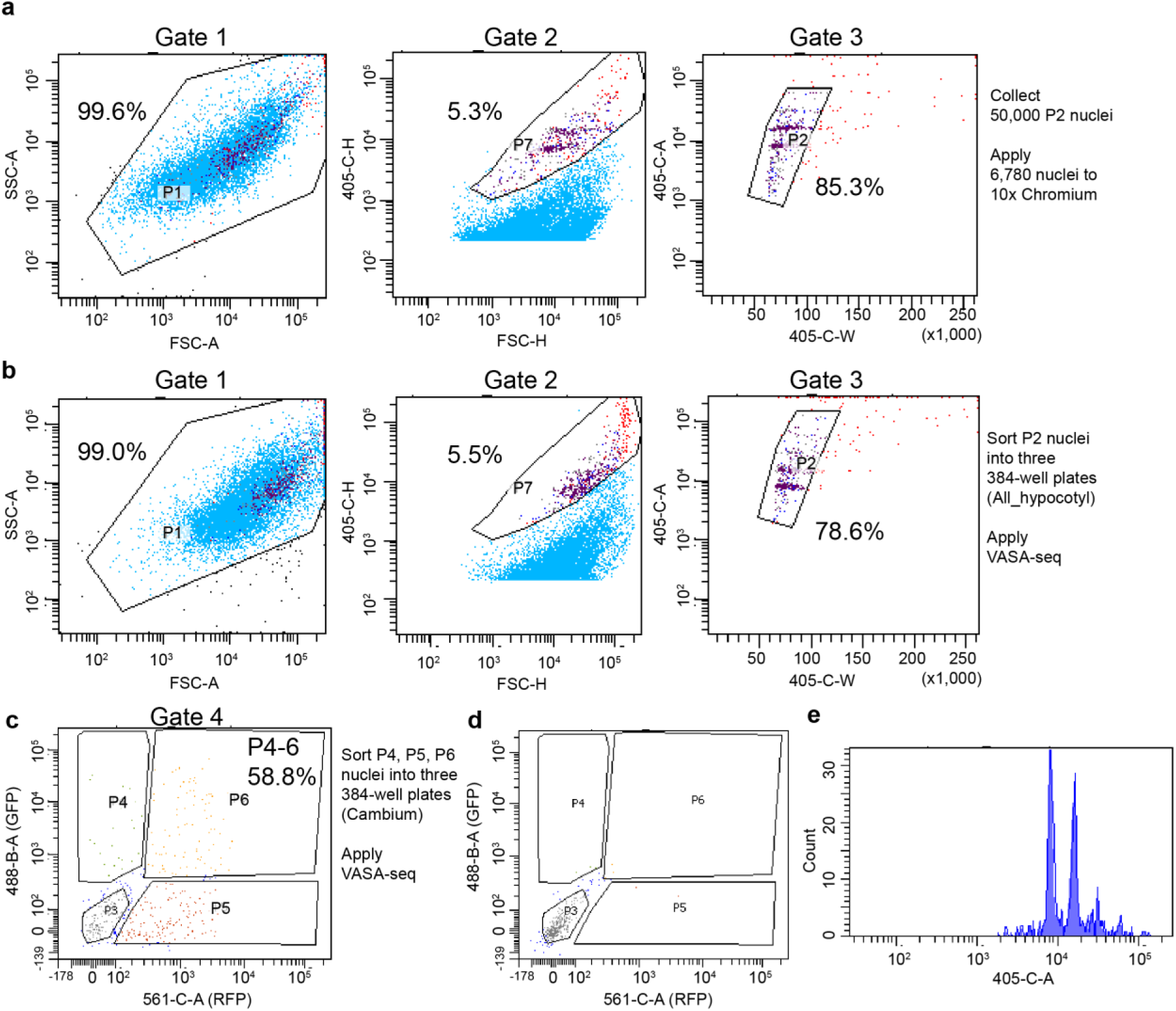
FACS gate settings for nucleus purification. **a**, Gate setting and dot plot of the FACSAria cell sorter for nucleus purification and 10× Chromium application. Nuclei were purified through Gate P1, P7 and P2 in a sequential manner. The percentages of each population compared to the parent population are indicated. FSC: forward scatter; SSC: side scatter; 405-C: fluorescence excited by 405nm laser (hoechst nucleus staining); -A:area; -H: hight; -W: width. **b–c**, Gate setting and dot plot of the FACSAria cell sorter for nucleus purification and VASA-seq application. Nuclei were purified through Gate P1, P7 and P2 in a sequential manner. Individual P2 nuclei were sorted into single wells of three 384-well plates as ‘nuclei from the whole hypocotyl’ (**b**). **(c)** P2 nuclei were further gated by fluorescence intensity, and individual GFP-positive, RFP-negative (P4), GFP-negative, RFP-positive (P5), or GFP-positive, RFP-positive (P6), nuclei were collected into single well of three 384-well plates as ‘cambium nuclei’. *PXY_pro_:H4-GFP*;*SMXL5_pro_:H2B-RFP* plants were prepared independently for (**b)** and (**c)**. **d,** Dot plot of the FACSAria cell sorter obtained from wild type plants using the same gate settings as used in (**c**) as a control. **e,** A histogram of hoechst signals of P2 nuclei demonstrating nucleus integrity.

**Supplementary Fig. S2:**
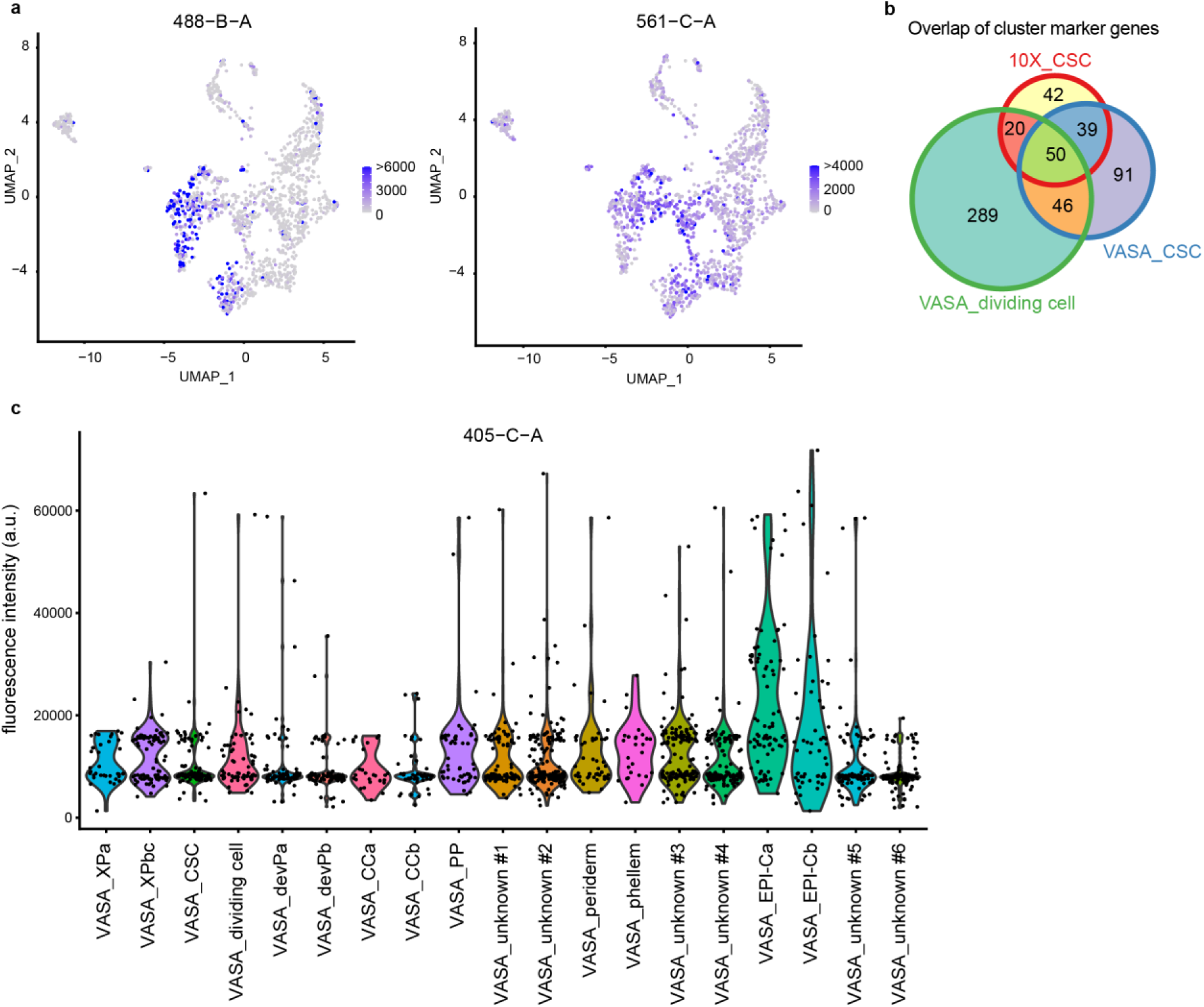
Nucleus fluorescence detected during VASA-seq analyses and CSC-specific marker genes. **a**, UMAP plots showing the fluorescence intensity of nuclei collected from *PXY_pro_:H4-GFP*;*SMXL5_pro_:H2B-RFP* plants captured during the sorting and excited by 488 nm (GFP, left) or 561 nm (RFP, right) laser light, respectively. **b**, Venn diagram showing the overlap of cluster-specific marker genes identified in the 10X_CSC, VASA_CSC and VASA_dividing cell clusters. **c**, Violin plot showing the Hoechst nuclear fluorescence signal excited by 405 nm laser light for each nucleus cluster.

**Supplementary Fig. S3:**
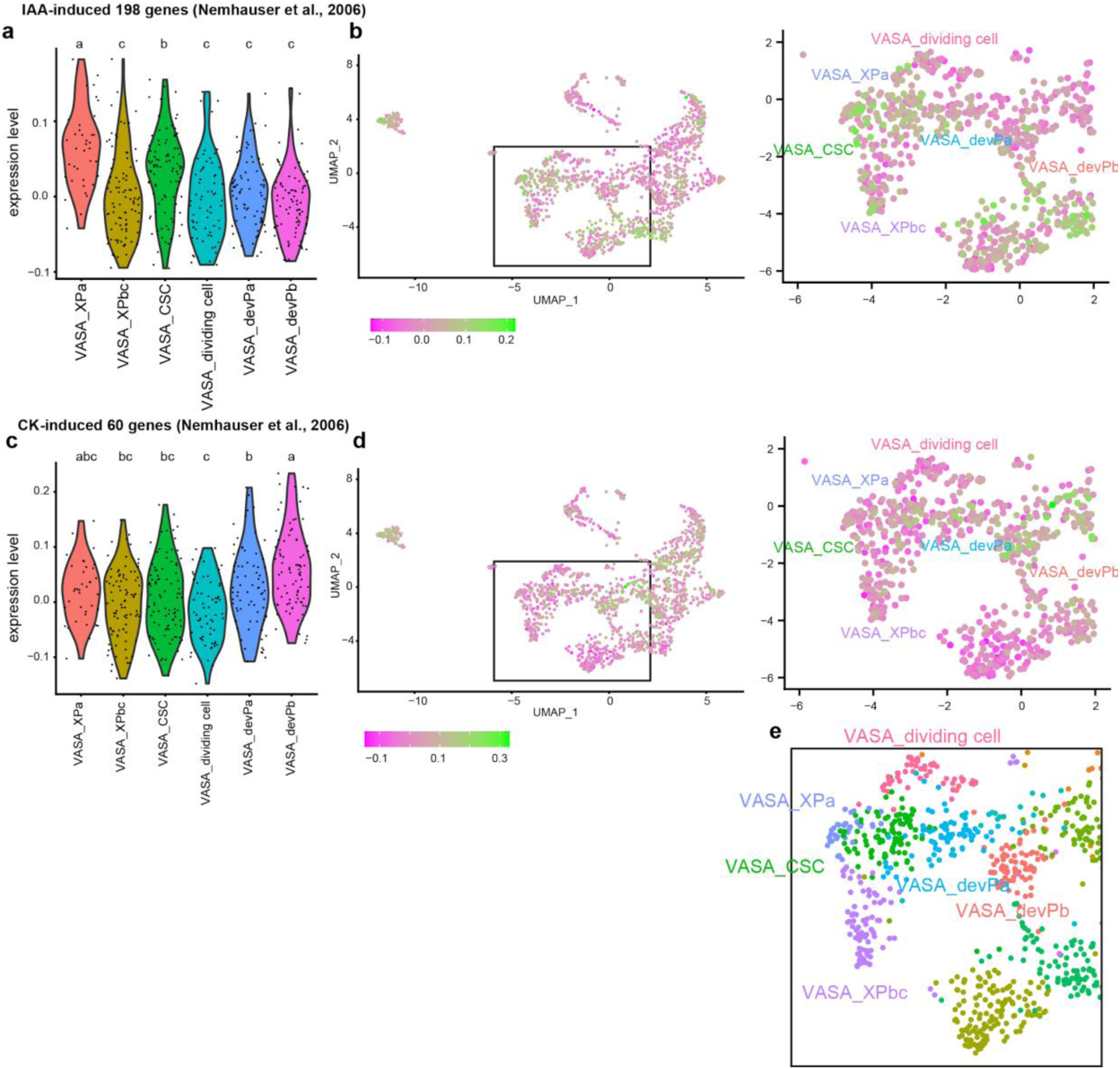
Expression of indole-3-acetic acid (IAA; auxin) inducible genes and zeatin (CK; cytokinin) inducible genes as detected during VASA-seq analyses. **a–d**, Violin plot (**a, c**) and UMAP visualisation (**b, d**) of transcript abundance of 198 IAA-inducible genes^26^ (a, b) and 60 CK-inducible genes^26^ (**c, d**) in the cambium-related cell clusters. The result of the Steel-Dwass test for multiple comparisons is indicated by letters (*p* < 0.05). Gene lists used in this analysis can be found at Supplementary Data 3. e, A close-up of the UMAP generated during VASA-seq cluster identification (Fig. 2) is shown as a reference.

**Supplementary Fig. S4:**
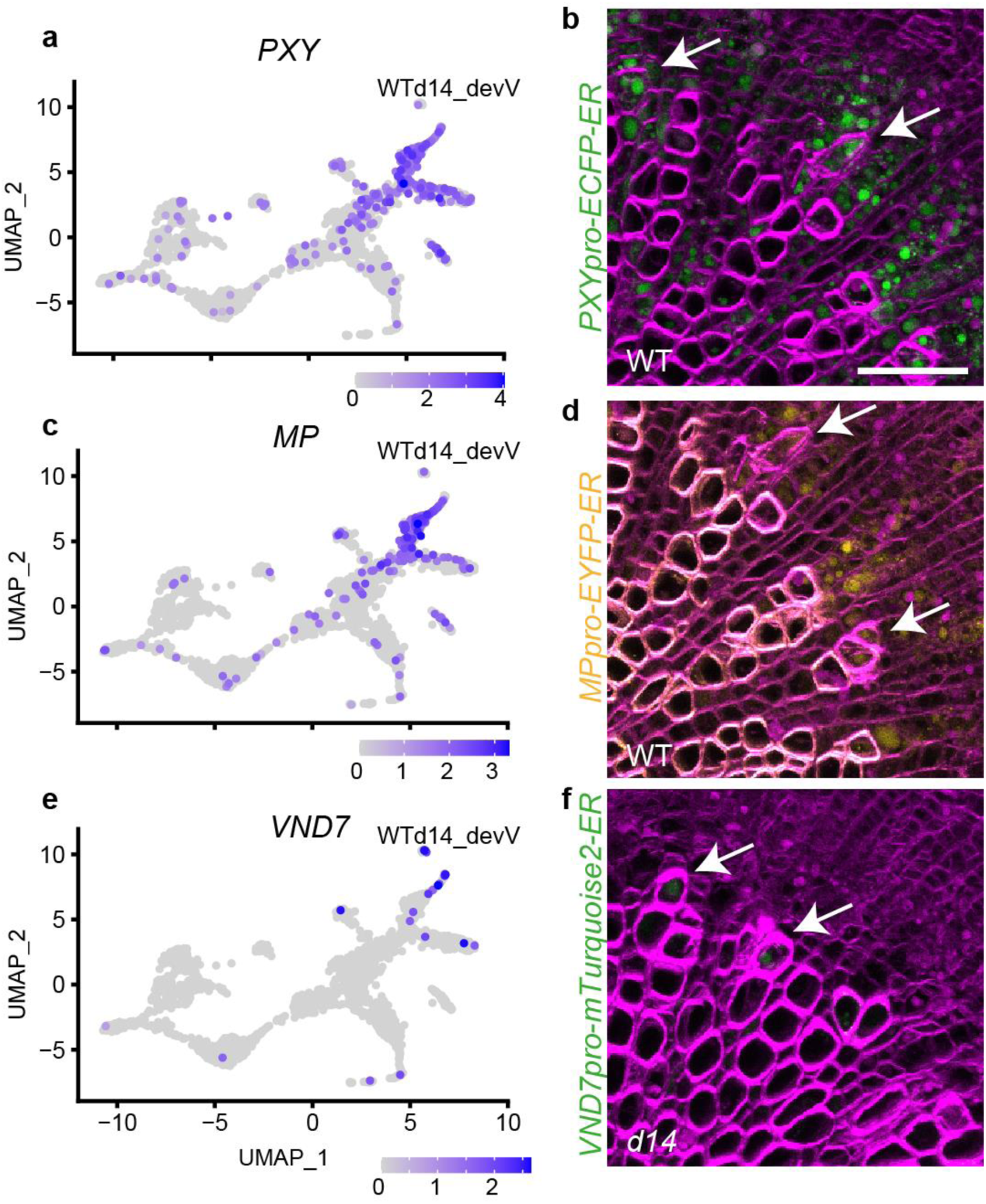
Expression pattern of *PXY*, *MP*, and *VND* in the UMAP plot and in the promoter reporter transgenic lines. **a, c, e**, Transcript abundance of the *PXY*, *MP* and *VND7* genes in the UMAP plot shown in Fig. 5. **b, d, f**, Hypocotyl cross-sections from plants carrying *PXY_pro_:ECFP-ER* (**e**), *MP_pro_:EYFP-ER* (**g**) or *VND7_pro_:mTurqouise2-ER* transgenes (e, g: wild type; i: *d14*) (**i**). Arrows indicate developing vessel elements. Scale bar represents 50 μm.

**Supplementary Fig. S5:**
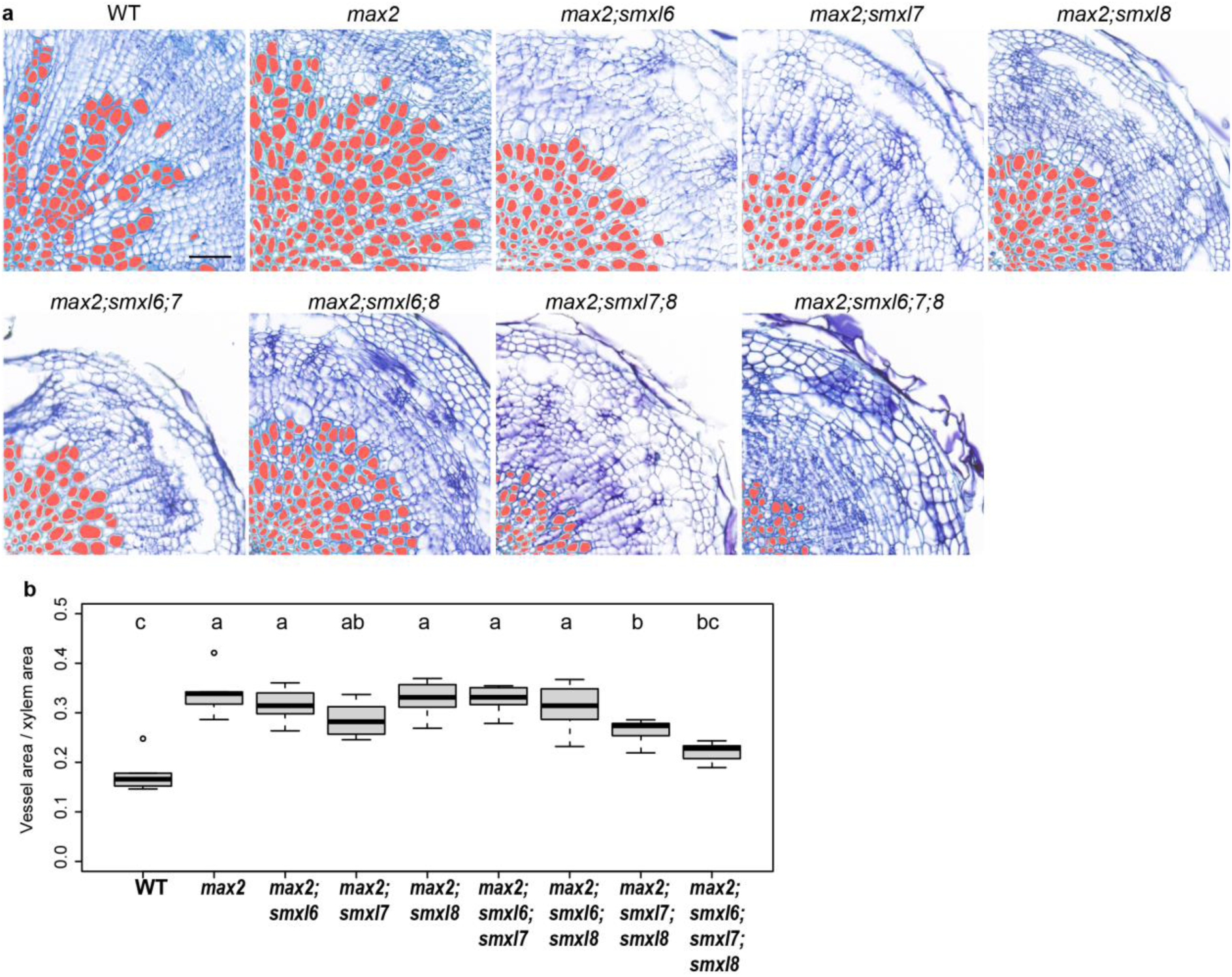
Histological analysis of higher order *max2*, *smxl6*, *smxl7* and *smxl8* mutants. **a**, Toluidine blue-stained hypocotyl cross-sections from five week-old wild type (WT), *max2* and higher order mutants in various combinations. Scale bar represents 50 μm. **b**, Quantification of the ratio between the vessel element area and the total xylem area in each mutant. n=5-13 plants for each genotype. Statistical groups are indicated by letters and were determined by a one-way ANOVA with post-hoc Tukey-HSD (95 % CI).

**Supplementary Fig. S6:**
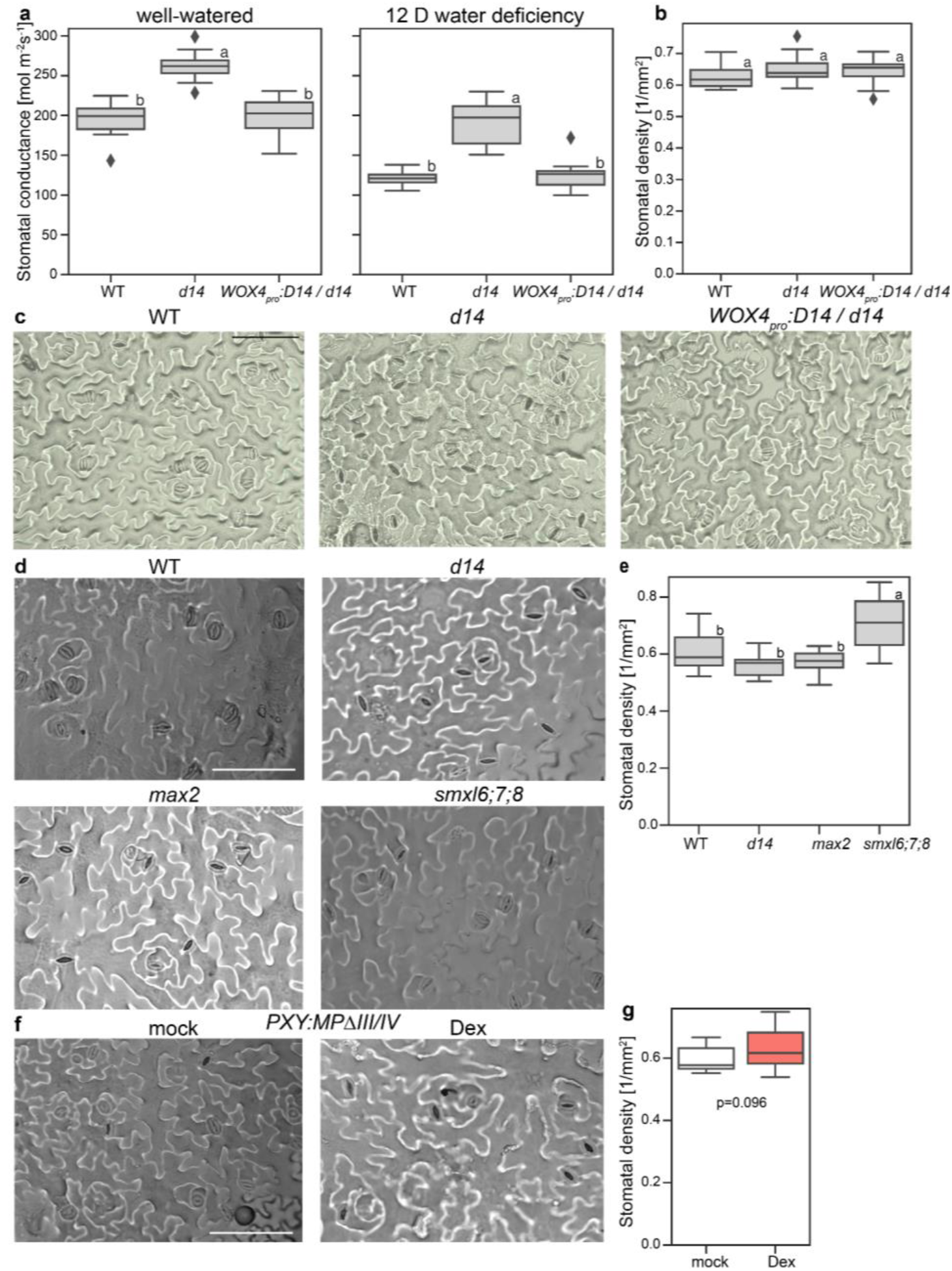
Stomata density analysis in SL-signalling mutants and upon auxin signalling modulation in vascular tissues. **a**, Quantification and comparison of stomatal conductance found in wild type, *d14*, and *WOX4pro:D14;d14* plants under well-watered conditions and 12 days (12 D) after initiating water deficiency treatments (n=17-19). Conductance was measured on three leaves per plant. Plots show average conductance of the three leaves per plant. **b**, Stomatal density (stomata per mm^2^) comparing genotypes shown in (**c**). n=17-19 plants for each genotype obtained from three independent experiments. **c**, Photomicrographs of the abaxial leaf surface of the same plants used for the 12 D water deficiency stomatal conductance measurement of wild type, *d14*, and *WOX4pro:D14;d14* plants. Size bar indicates 50 μm. **d**, Photomicrographs of the abaxial leaf surface of wild type, *d14*, *max2*, and *smxl6;7;8*. Same plants used for stomatal density analysis as for the 12 D water deficiency stomatal conductance measurement. Size bar indicates 50 μm. **e**, Stomatal density (stomata per mm^2^) comparing genotypes shown in (**d**). n=15-20 plants for each genotype obtained from three independent experiments. **f-g**, Photomicrographs and stomatal density on the abaxial leaf surface of mock-or Dex-treated plants carrying a *PXY_pro_:GR-MPΔIII/IV* transgene. n=14 (mock), 16 (Dex) plants obtained from three independent experiments. Statistical groups are indicated by letters and were determined by a one-way ANOVA with post-hoc Tukey-HSD (95% CI). Size bar indicates 50 μm.

**Supplementary Table 1:**
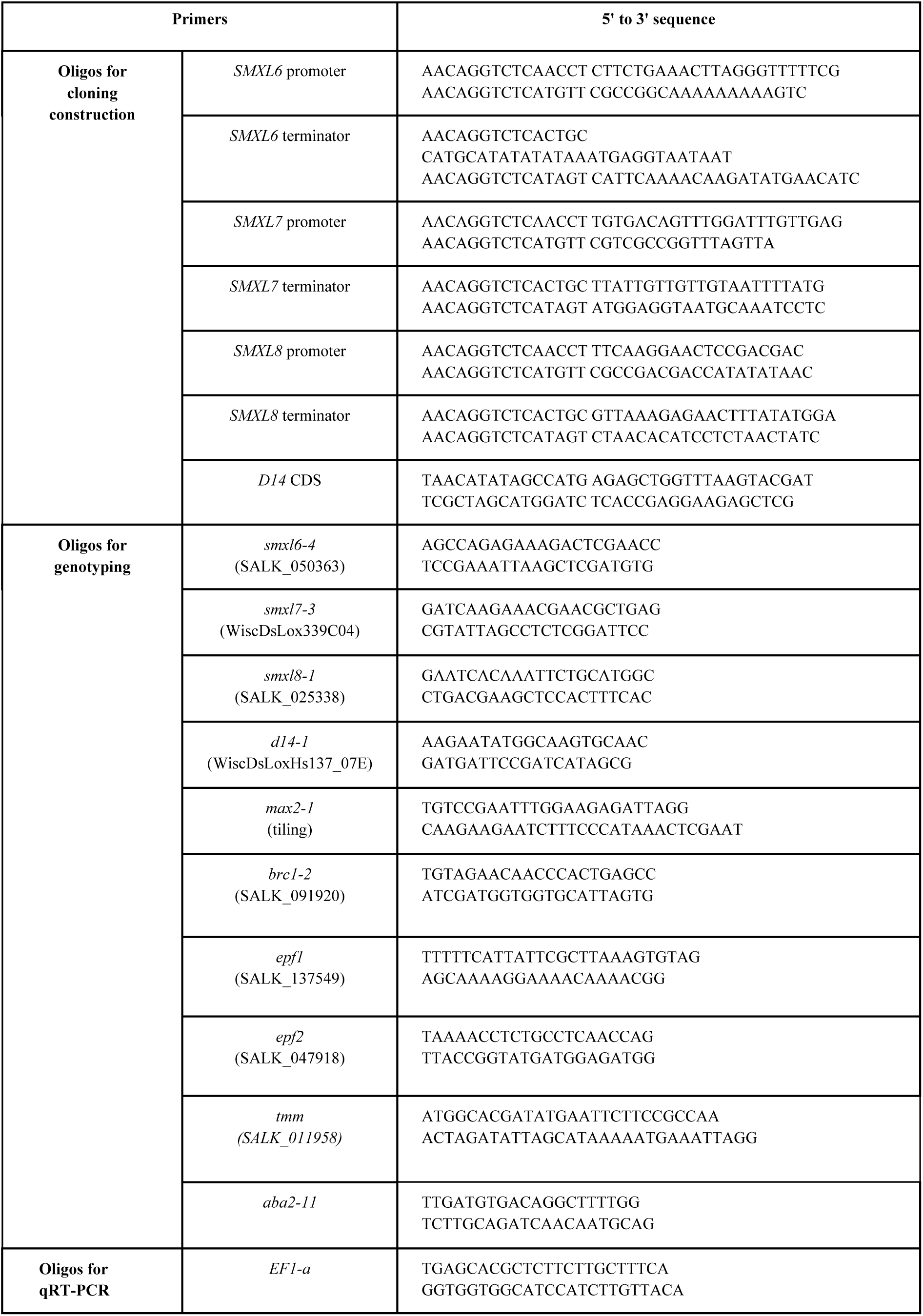
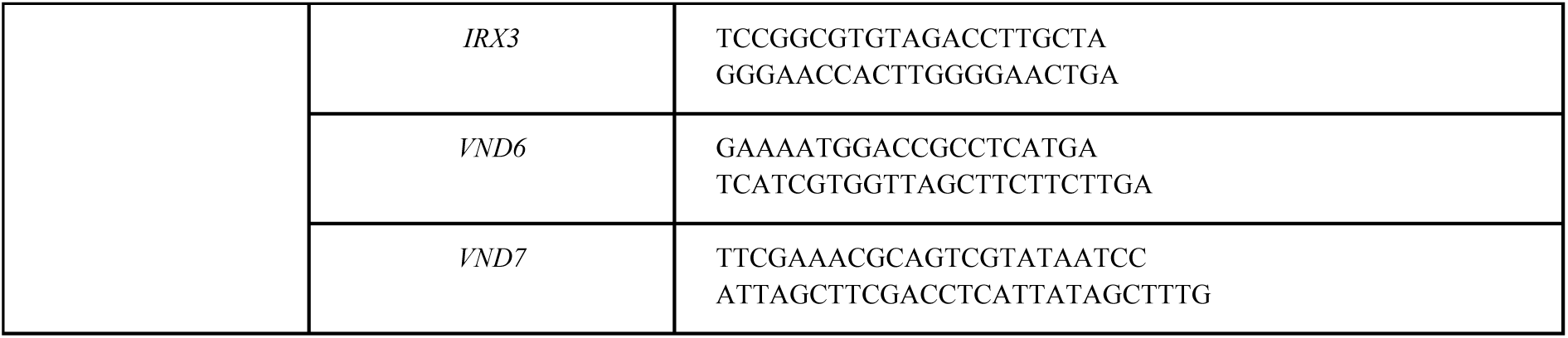
Oligo sequences used in this study.

**Supplementary Table 2:**
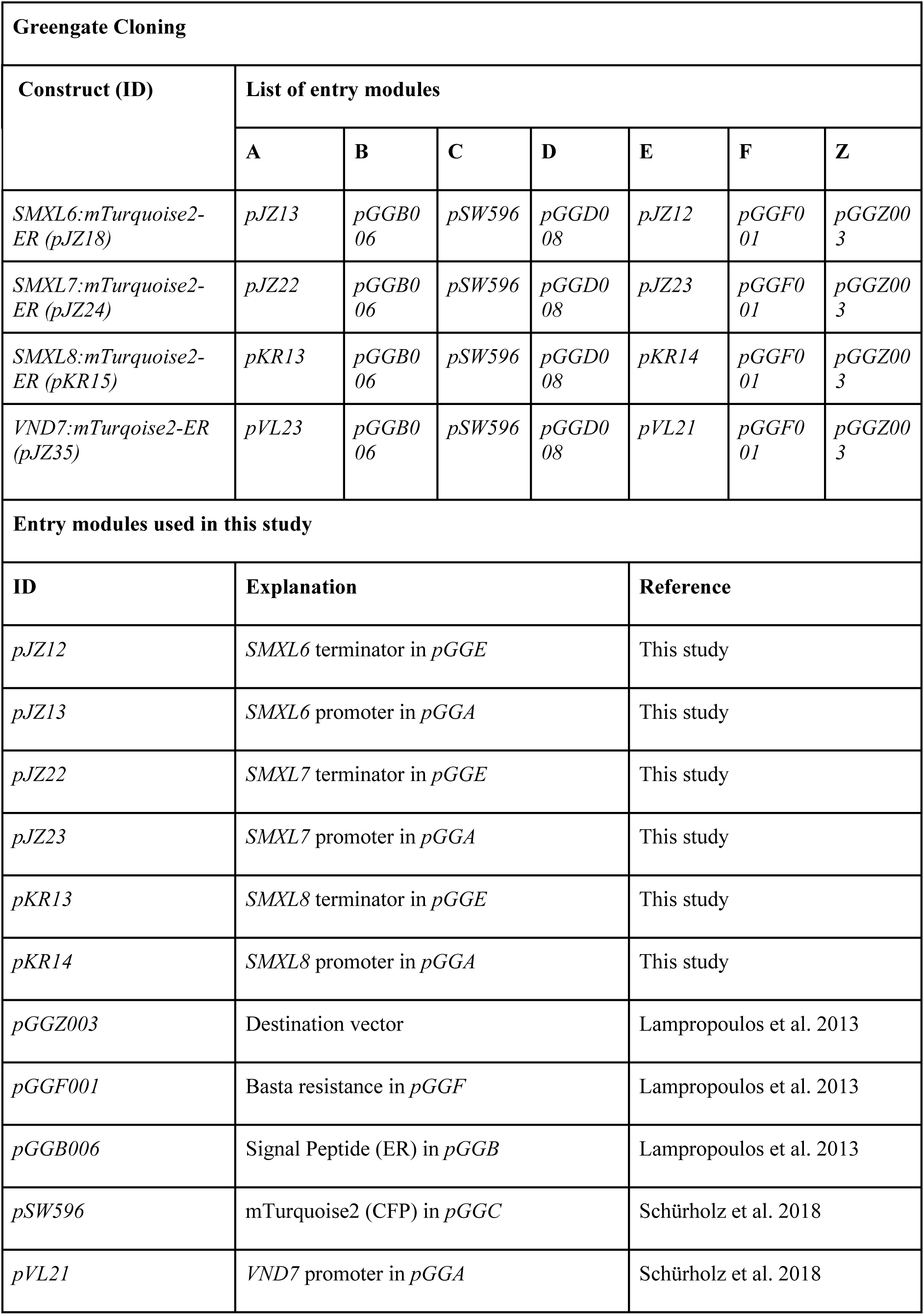
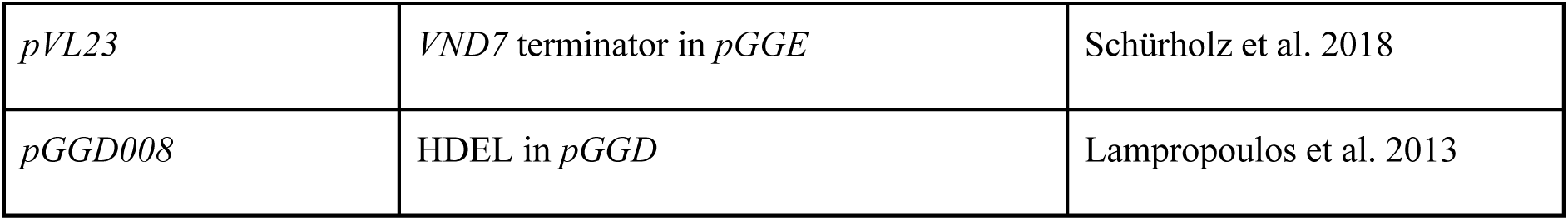
Vector construction strategy used in this study.

| **Greengate Cloning** |  |  |  |  |  |  |  |
| --- | --- | --- | --- | --- | --- | --- | --- |
| **Construct (ID)** | **List of entry modules** |  |  |  |  |  |  |
|  | **A** | **B** | **C** | **D** | **E** | **F** | **Z** |
| *SMXL6:mTurquoise2-ER (pJZ18)* | *pJZ13* | *pGGB006* | *pSW596* | *pGGD008* | *pJZ12* | *pGGF001* | *pGGZ003* |
| *SMXL7:mTurquoise2-ER (pJZ24)* | *pJZ22* | *pGGB006* | *pSW596* | *pGGD008* | *pJZ23* | *pGGF001* | *pGGZ003* |
| *SMXL8:mTurquoise2-ER (pKR15)* | *pKR13* | *pGGB006* | *pSW596* | *pGGD008* | *pKR14* | *pGGF001* | *pGGZ003* |
| *VND7:mTurquoise2-ER (pJZ35)* | *pVL23* | *pGGB006* | *pSW596* | *pGGD008* | *pVL21* | *pGGF001* | *pGGZ003* |
| **Entry modules used in this study** |  |  |  |  |  |  |  |
| **ID** | **Explanation** |  |  |  | **Reference** |  |  |
| *pJZ12* | *SMXL6* terminator in *pGGE* |  |  |  | This study |  |  |
| *pJZ13* | *SMXL6* promoter in *pGGA* |  |  |  | This study |  |  |
| *pJZ22* | *SMXL7* terminator in *pGGE* |  |  |  | This study |  |  |
| *pJZ23* | *SMXL7* promoter in *pGGA* |  |  |  | This study |  |  |
| *pKR13* | *SMXL8* terminator in *pGGE* |  |  |  | This study |  |  |
| *pKR14* | *SMXL8* promoter in *pGGA* |  |  |  | This study |  |  |
| *pGGZ003* | Destination vector |  |  |  | Lampropoulos et al. 2013 |  |  |
| *pGGF001* | Basta resistance in *pGGF* |  |  |  | Lampropoulos et al. 2013 |  |  |
| *pGGB006* | Signal Peptide (ER) in *pGGB* |  |  |  | Lampropoulos et al. 2013 |  |  |
| *pSW596* | mTurquoise2 (CFP) in *pGGC* |  |  |  | Schürholz et al. 2018 |  |  |
| *pVL21* | *VND7* promoter in *pGGA* |  |  |  | Schürholz et al. 2018 |  |  |

|  |  |  |
| --- | --- | --- |
| *pVL23* | *VND7* terminator in *pGGE* | Schürholz et al. 2018 |
| *pGGD008* | HDEL in *pGGD* | Lampropoulos et al. 2013 |

## Other Supplementary Items

### Methods description

Supplementary Data 1

Basic statistics of single nucleus (sn) RNA-seq analyses presented in this study.

Supplementary Data 2

Annotation and marker genes detected for each cluster in all the snRNA-seq analyses presented in this study.

Supplementary Data 3

Gene lists used in snRNA-seq analyses obtained from previous studies and current study.

Supplementary Data 4

Fluorescent signal of each nucleus for each multiwell plate obtained during FANS for VASA-seq analysis.

Supplementary Data 5

Zip-archived Seurat object files (10×_All_hypocotyl.rds, VASA-seq.rds and WTd14.rds) are provided for retrieving gene expression patterns in each snRNA-seq analysis dataset. Remarks are mentioned in Supplementary Data 2.

Supplementary Data 6

*p* values obtained in t-test at each timepoint between each mutant genotype and wildtype in pot weight measurements used in Fig. 9a.

