## Supplementary material for "Strigolactones optimise plant water usage by modulating vessel formation": Methods

### Plant material

All plant lines used in this study were *Arabidopsis thaliana* (L.) Heynh. plants of the accession Columbia (Col-0). The single and higher order mutants of *d14-1* (WiscDsLoxHs137_07E), *smxl6-4* (SALK_049115), *smxl7-3* (WiDsLox339_C04), *smxl8-1* (SALK_025338C) and *max2-1* were obtained from Dave Nelson[1,2] (UC Riverside, US) and Ottoline Leyser[3] (SLCU, Cambridge, UK). The *brc1-2* (SALK_091920) mutant was ordered from NASC and was characterised previously[4–7]. *Strigo-D2*[8], $PXY_{pro}$*:ER-ECFP-HDEL*[9], $PXY_{pro}$*:H4-GFP;*$SMXL5_{pro}$*:H2B-RFP*[10], $SMXL7_{pro}$:$SMXL7^{d53}$*-VENUS*[11], $PXY_{pro}$*:Myc-GR-bdl*[12], $MP_{pro}$*:ER-EYFP-HDEL*[12] and $PXY_{pro}$*:GR-MPΔIII/IV*[12] transgenic lines were described previously. Other transgenic lines were generated through the floral dipping method using *Agrobacterium tumefaciens*[13]. The *aba2-11* mutant[14] was obtained from Mikael Brosché (University of Helsinki, Finland). *epf1-1* (SALK_137549)[15] and *epf2-3* (SALK_047918)[16] single and double mutants and *tmm-1* (SALK_011958)[17] mutants were donated by Christopher Grefen (University of Bochum, Germany).

### Vector construction

The $WOX4_{pro}$*:D14* (*pVJ13*) construct was generated using In-Fusion cloning (Takara Bio) using the amplified *D14* coding region and BamHI-digested *pTOM49*[18]. Other plasmids were generated using the GreenGate cloning system[19,20]. See Supplementary Table 1 for oligo sequences used for molecular cloning and Supplementary Table 2 for each module used during the cloning process.

**Growth conditions**

Arabidopsis seeds were surface sterilised by 70 % ethanol containing 0.02 % Tween-20, stratified at 4°C for 2-3 days in the dark, sown on 0.8 % w/v agar in 1/2 Murashige and Skoog (MS) medium supplemented with 1% w/v sucrose. Seedlings were grown in short-day conditions (SD; 10 h light and 14 h darkness) at 21-22°C. For morphological observations and reporter activities analyses, 5 day-old seedlings were transferred to pots filled with 4:1 mixture of soil and vermiculite. After 21 days, plants were transferred to long day conditions (LD; 16 h light and 8 h darkness) at 21-22°C. Plants for stomatal conductance measurements were grown for six weeks in SD conditions (8.5 h light and 15.5 h darkness).

**Single nucleus RNA-seq analysis**

Hypocotyls were dissected and collected in petri dishes incubated on ice. 2 ml of 1x nuclei isolation buffer (CelLytic ™ PN Isolation/Extraction Kit, Sigma #CELLYTPN1) supplemented with 20 µl RiboLock RNase inhibitor 40 U/µL (ThermoFisher #EO0381) and Hoechst 33342 at 10 µg/ml final concentration were prepared and a minimum amount of buffer was applied to submerge the collected hypocotyls[10]. Hypocotyls were chopped using razor blades (Wilkinson Sword) for up to 5 min and transferred on a gentle shaker at 4°C for 15 min. Samples were then filtered through a 50 µm filter (CellTrics, Sysmex #04-004-2327) and passed to a low protein binding tube (Eppendorf #0030108132).

For 10x genomics application, 50,000 nuclei were sorted into 33 µl collection buffer (10 µl PBS (Corning, #21-040-CV), 5 µl BSA (ThermoFisher, #AM2616), 6 µl RNAse inhibitor (ThermoFisher, #AM2682), 12 µl RNAse inhibitor (ThermoFisher, #AM2694)), modified from a previous report[21], by a BD FACSAria IIIu cell sorter (Becton Dickinson) using a 100-µm sort nozzle (see Supplementary Fig. S1). A sheath pressure of 35 psi and a drop drive frequency of 60 kHz were applied. 43.2 µl of sorted nuclei solution was applied to a Chromium Next GEM Chip without dilution and Chromium Next GEM Single Cell 3' GEM, Library & Gel Bead Kit (v3.1) was used to generate libraries following the manufacturer's instruction. Nuclei concentration was monitored by a fluorescent cell counter (ThermoFisher, Countess 3 FL) using the DAPI channel to obtain approximately 200 nuclei/µl. Libraries were sequenced using a NextSeq (Illumina) machine at the high output mode.

For initial wild type analyses, about 100 hypocotyls from a *PXY*$_{pro}$*:H4-GFP;SMXL5*$_{pro}$*:H2B-RFP* line collected four weeks after germination were used. Sequencing reads were mapped to the Arabidopsis genome (with H4GFP and H2BRFP transgene sequences added) using STAR (2.7.8a)[22] with the "--alignIntronMax 10000 --alignMatesGapMax 10000" option. "GeneFull" option was used in STAR solo to include reads mapped to introns. Seurat[23] (4.0.6) was used for further analysis. Cells with nCount_RNA between 1201 and 9999 and a mitochondrial genome read fraction of less than 20 % were kept for further analysis. Clustering and UMAP were generated using default settings of Seurat with the parameters „dims = 1:15, resolution =1.2, algorithm =2".

For wild type and *d14* mutant analysis, about 200 hypocotyls collected 19 days after germination were used. Sequencing reads were mapped to the Arabidopsis nuclear genome with cell ranger (6.0.1, 10x genomics) using the "--include-introns" and "--alignIntronMax 10000 --alignMatesGapMax 10000" alignment options. nCount_RNA between 1501 and 14999 were kept for comparing wild type and *d14* mutants. Clustering was carried out using parameters „dims = 1:15, resolution =1.2, algorithm =2", and UMAPs were generated after randomly resampling 500 nuclei from each genotype.

For VASA-seq analysis[24], about 100 hypocotyls from a *PXY*$_{pro}$*:H4-GFP;SMXL5*$_{pro}$*:H2B-RFP* line were collected four weeks after germination to obtain 1,134 'cambium region' nuclei (three 384-well plates). Additional 100 hypocotyls were used for collecting 1,134 'all region' nuclei (three 384-well plates) (see Supplementary Fig. S1). Single nuclei were sorted into individual wells of 384-well plates containing well index oligos purchased from Single Cell Discoveries B.V. (Utrecht, The Netherlands), using the index sorting mode of the BD FACSAria IIIu cell sorter, recording the fluorescence signal of each nucleus (Supplementary Fig. S2a, c, Supplementary Data 4). Multi-well plates were frozen and further processed according to the VASA-seq protocol[24] of Single Cell Discoveries with the following modifications. 1) In the end repair and polyA reactions, our added mix per well contained 7.5 µM ATP and 3,75 mU of polyA polymerase. 2). We used 6 µl of ExoSAP after *in vitro* transcription. 3) The composition of our 2,5x RNaseH buffer used during the rRNA depletion step was 125 mM Tris-HCl pH 7.5, 250 mM NaCl, 10 mM $MgCl_2$. Sequencing reads were similarly mapped to the Arabidopsis genome (with H4GFP and H2BRFP transgene sequences added) using STAR with the same options mentioned above. Well barcodes were obtained from Gene Expression Omnibus (https://www.ncbi.nlm.nih.gov/geo/; Accession ID: GSE112438, celseq2_bc.csv.gz). Fluorescent signals were combined in Seurat with a 150 offset value to avoid negative values in the 488 nm and 561 nm channels and all nuclei data were merged. Cells with nCount_RNA values between 1201 and 14999, a mitochondrial genome read fraction lower than 20%, fluorescent signals in the 405 nm and 488 nm channels lower than 75,000 were kept for further analysis. Clustering and UMAP were performed with parameters „dims = 1:15, resolution =1.8, algorithm =2".

Basic statistics of all single cell analyses can be found in Supplementary Data 1. Marker genes for each cluster were generated by using the *FindAllMarkers* function in Seurat (Supplementary Data 2) (only.pos = TRUE, min.pct = 0.25, logfc.threshold = 0.25, test ="wilcox", return.thresh = 0.01). Seurat object files for each dataset can be found in Supplementary Data 5. Phytohormone responsive genes[25,26] and marker genes for cell clusters generated by single cell transcriptome analyses[27,28] were curated from previous reports and integrated in Seurat by using the AddModuleScore function (Supplementary Data 3).

Differential expression was tested by the Steel-Dwass test using an R script (http://aoki2.si.gunma-u.ac.jp/R/src/Steel-Dwass.R) authored by Shigenobu Aoki.

**Confocal Microscopy**

Hypocotyl samples were fixed overnight at 4°C in 4 % (w/v) PFA dissolved in PBS. The tissue was washed twice with PBS, embedded in 5 % low melting agarose, sectioned by razor blades (Wilkinson basic), and then stained with 0.1% DirectRed 23 or 10 μg/ml Hoechst 33342 for 5 minutes at room temperature. Excess staining was removed by clearing the sample in 1X PBS. Confocal microscopy experiments were carried out on a Leica TCS SP8 (Leica Microsystems; Mannheim, Germany). 458 nm, 514 nm and 561 nm lasers were used to excite mTurquoise2 (CFP), YFP (mVenus), and mCherry/Direct Red, and emissions were detected at 465-509 nm, and 524-540 nm and 571-630 nm, respectively. Hoechst 33342 and Calcofluor White, together with lignin in differentiated xylem vessel elements were visualised using a 405 nm laser, and collection of the emission at 410-450 nm.

**Strigo-D2 ratio analysis**

False colour images were generated using ImageJ through calculating intensity ratios of each pixel from mVenus and mCherry channels after being Gaussian Blurred and subtracting background signals. For calculating the ratio value of each nucleus, nuclear regions were detected by using the Particle Analyzer function in ImageJ after masking the nuclear region through thresholding. Then, nuclear mVenus and mCherry signal intensity were measured and intensity ratios were determined. Nuclei within cambium, phloem, and xylem zones, as well as in developing vessel elements were manually defined as follow: around six cell layers counting from the vessel element border toward the organ periphery were defined as cambium zone.

Nuclei distal to the cambium were defined as phloem and the nuclei proximal to the cambium were defined as xylem. The enlarged nuclei within the xylem region were considered as being located in developing vessel elements.

**Histological analyses**

The harvested hypocotyls from five week-old (3 weeks SD + 2 weeks LD) plants were infiltrated in 70 % ethanol for at least three days at 4°C before being paraffin embedded by the Leica ASP200 S processor (Leica Microsystems, Mannheim, Germany). After embedding in paraffin, the microtome RM2235 (Leica Microsystems, Mannheim, Germany) was used to produce 10-µm thick sections. The sections were harvested from the upper part of the hypocotyl, 1 mm below the leaf primordia. Dried sections were deparaffinized, stained with 0.05 % toluidine blue (#52040, AppliChem, Darmstadt, Germany) and fixed by Micromount Mounting Media (Leica) on microscope slides (Thermo Scientific; Wal-tham, USA). Slides were scanned using the Pannoramic SCAN II scanner (3DHistech, Budapest, Hungary) and analysed by the CaseViewer 2.2 software (3DHistech, Budapest, Hungary). Vessel elements were automatically detected using the Particle Analyzer option in Fiji and adjusted manually to avoid false positives.

**Dexamethasone treatment**

Stock solution of 25 mM Dex was dissolved in DMSO, and a 15 µM working solution was freshly prepared by diluting the stock solution with tap water. Control treatments contained an equivalent amount of solvent. Plants were initially grown in SD conditions for three weeks without treatment, and treatment was started when plants were transferred to LD conditions by watering twice a week with either 50 ml 15 µM Dex or mock solution per pot until harvest.

**GR24$^{4DO}$ application**

GR24$^{4DO}$ (10 μM) was prepared through a 1000x dilution of a stock solution (10 mM GR24$^{4DO}$ dissolved in acetone). Seedlings were initially grown on MS medium plates for two weeks (SD conditions) without treatment, subsequently transferred to plastic containers supplemented either with 10 μM GR24$^{4DO}$ or mock solution containing an equivalent amount of acetone, grown for another two weeks (1 week SD + 1 week LD) and harvested for histological analyses.

**qRT-PCR analysis**

Hypocotyls of four week-old wild type and *d14* mutant plants grown in soil were harvested. Total RNA was extracted using a RNeasy Mini Kit (Qiagen, Hilden, Germany), followed by genomic DNA digestion according to the TURBO DNA-free™ Kit (Thermo Scientific; Wal-tham, USA) protocol. cDNA synthesis was performed according to the instructions of the Thermo Revert Aid Kit (Thermo Scientific; Wal-tham, USA). Real-time PCR assays were conducted using SYBR Green Mix (Thermo Scientific; Wal-tham, USA) and gene-specific primers (Supplementary Table 1) on a qTOWER3 thermal cycler with using the *EF1-a* (*AT5G60390*) gene as an internal reference.

**Pot weight measurements during water deficiency treatments**

To grow plants under comparable conditions, each pot contained 70 g of soil. For watering, pots were placed in petri dishes to soak 25 ml water overnight from below. Next, 20 ml of nematode-containing solution was added to each pot from above. Plants were kept in petri dishes during subsequent watering (20 ml every week during the first three weeks and 25 ml twice a week for the last 2 weeks). Pot weight was measured starting at five weeks after germination for consecutive 12 days.

**Stomata conductance measurements**

Wild type, *d14, max2* and *smxl6;7;8* plants were grown in growth chambers with sufficient watering for 6.5 weeks, followed by water deficiency treatments for 12 days. During the first 6.5 weeks, plants were watered with 20 ml every week during the first three weeks and 30 ml twice a week for the next 3.5 weeks. Dex or mock-treated *PXY$_{pro}$:MPΔIII/IV* plants were not watered for 10 Days after the Dex or DMSO treatment. Stomata conductance of wild type, *d14*, *max2* and *smxl6;7;8* plants was measured after 12 days of water deficiency treatment and Dex or mock-treated *PXY$_{pro}$:MPΔIII/IV* plants were analysed after 10 days of water deficiency treatments. For the measurements, the SC-1 Leaf Porometer (METER Environment ®) was clipped on the abaxial side of the leaf. The 9$^{th}$, 10$^{th}$ and 11$^{th}$ produced leaf of wild type, *smxl6;7;8,* Dex or mock-treated *PXY$_{pro}$:MPΔIII/IV* plants and three well expanded upper leaves from *d14* or *max2* mutants were chosen for the measurement. The SC-1 Leaf Porometer (Meter Group, Pullman, US) was used according to the user manual. Single measurements were temporarily randomised across the course of the day and across genotypes.

**Stomata Imprints**

Water deficiency-treated plants used for the stomatal conductance measurement were taken to imprint the abaxial leaf side for stomata density quantification (stomata/mm$^2$). For the imprint, a small drop of instant adhesive glue UHU Sekundenkleber blitzschnell (UHU, Bühl, Germany) was placed on a Superfrost™ Microscope slide and the leaf was gently pressed on the glue for two seconds. The imprints were visualised using a Contrast Microscope DMIRB microscope with a 20x objective and the bright field mode. Five images of the central leaf regions of each imprint were taken. The number of stomata was counted using Fiji.

**Basic Fuchsin staining**

To observe xylem strands in roots, five day-old seedlings were stained and fixed in 0.2 % (m/v) Basic Fuchsin dissolved in ClearSee[29,30] (10 % xylitol, 15% sodium deoxycholate, and 25 % UREA) solution overnight. Next the fuchsin solution was removed and samples were washed once with ClearSee for 30 min. Subsequently, seedlings were stored in ClearSee solution and analysed.

**Statistics**

All the measurements were done on distinct samples. Statistical tests were applied in a two-sided mode. Box plots indicate the 25th (Q1, box limit), 50th (median, centre line) and 75th (Q3, box limit) percentiles and whiskers indicate the value range or up to the 1.5x interquartile range from the Q1 or Q3 limit, respectively. Data points beyond this range were plotted individually as outliers. Statistic analyses were carried out using R (v4.0.4), ggplot2 (v3.3.3), or Python (v3.10.7), pandas (v1.5.0) and seaborne (v0.12.0).
